# Elevated temperature drives the biosynthesis of novel acylated glucosinolates in *Arabidopsis thaliana* seeds

**DOI:** 10.64898/2026.06.03.729804

**Authors:** Léa Barreda, Stéphanie Boutet, Marine Lebon-Navarro, Nicolas Klewko, Céline Brosse, Elettra Frassineti, Céleste Angibaud, Ikram Bendraoua, Audrey Le Cabec, Delphine De Vos, Céline Boulard, Damaris Grain, Sébastien Baud, Loïc Rajjou, François Perreau, Loïc Lepiniec, Massimiliano Corso

## Abstract

Glucosinolates (GSLs) are major defensive compounds massively accumulated in Brassicaceae seeds, including that of the model plant *Arabidopsis thaliana*. While most studies have focused on the role of GSL in responses to biotic stress, the potential regulation and function of GSLs in responses to abiotic stresses have been neglected, particularly in seeds.

In this study, multi-omic analyses revealed a previously uncharacterized GSL modification pathway induced by elevated temperature (ET) during *A. thaliana* seed development. Activation of this pathway leads to the production of several novel thioglucose-acylated GSLs, including sinapoylated and benzoylated derivatives. A reverse genetics approach demonstrated that the SERINE CARBOXYPEPTIDASE-LIKE 17 (SCPL17) and BENZOYLOXYGLUCOSINOLATE 1 (BZO1) enzymes are required for the acylation of GSL thioglucose moieties. Furthermore, the accumulation of acylated GSLs in seeds of 85 *A. thaliana* accessions grown under standard condition was shown to correlate with the average annual temperature of their origin site, suggesting that thioglucose-acylated GSLs production may reflect long-term thermal adaptation across natural populations.

Taken together, these results demonstrate that thioglucose acylation by SCPL17 and BZO1 represents a new layer of GSL diversification in *A. thaliana* seeds that contributes to both ET response and long-term environmental adaptation.

## INTRODUCTION

Seed quality is a preeminent agricultural issue as it impacts both food and non-food applications, while also playing a central role in biodiversity preservation and environment protection. Seed quality is determined both by seed composition (e.g. specialized metabolites [SMs], storage oils, polysaccharides and proteins) and other intrinsic and extrinsic factors that impact seed ability to develop and germinate successfully in agriculture or natural environments. These traits are shaped by both genetic factors and environmental conditions (Hanif et al. 2023).

SMs are small organic compounds, often characterized by specific distributions across plant families, genera, or species. The large diversity of chemical structures, and consequently of biological functions, observed among metabolic categories arises from the addition of various functional groups to SM core structures. These modifications or decorations include hydroxylation, methylation, glycosylation and acylation (Barreda et al. 2024). SMs play crucial roles in the interactions of plants and seeds with their environment (Abbas et al. 2017; Corso et al. 2020). Moreover, they constitute high added-value molecules with a broad range of applications in food, health, medicine, cosmetics and agroecology industry (Corso et al. 2020; Baud et al. 2022; Barreda et al. 2024). SM synthesis and accumulation is strongly influenced by the environmental conditions.

Temperature is a major factor affecting growth, development and quality of plants and related products, including seeds. Indeed, agronomic crop production is critically affected by considerable increases of temperature and cause drastic seed yield losses. It has been reported that yields of the 4 major crops (wheat, rice, maize and soybean) are reduced by roughly 6.0%, 3.2%, 7.4%, and 3.1% on average, respectively for every 1°C increase in global temperature (Zhao et al. 2017). In Brassicaceae species such as *A. thaliana, Brassica napus* and *Camelina sativa*, seed embryo development is initially accelerated by ET but becomes affected following prolonged exposure, which can lead to seed abortion (Mácová et al. 2022; Nadakuduti et al. 2023; Barreda et al. 2025a). Furthermore, ET affects seed photosynthetic activity and the accumulation of storage compounds such as proteins, oils and other carbohydrates (Brunel-Muguet et al. 2015; Mácová et al. 2022; Nadakuduti et al. 2023).

The distribution and phenology of plant species across the world has already altered, due to climate change (Scheffers et al. 2016), highlighting that plants are able to develop mechanisms to respond and to adapt to temperature rise. Biological responses correspond to short-term reactions to a stimulus (e.g. rise of temperature), whereas adaptation relate to long-term, structural, or functional change that enhances survival of specie or individual over time (Zhang et al. 2022; Munns and Millar 2023). World global mean temperature is expected to increase of 0.3°C per decade and the occurrence and intensity of heat waves are predicted to increase in the coming years (Guihur et al. 2022), hence plants ET responses and thermal adaptation will pursue and possibly evolve.

Seeds of the model plant *A. thaliana* accumulate a wide range of SMs, including the Brassicaceae-specific nitrogen-containing glucosinolates (GSLs) and many phenylpropanoids (e.g. cinnamic acids, flavonols and proanthocyanidins) (Matsuda et al. 2010; Barreda et al. 2025a). Hence, *A. thaliana* constitutes a valuable model to study SM diversity and functions in Brassicaceae seeds. The synthesis of GSL core structures occurs in vegetative tissues and in a specialized reproductive structure, the funiculus, from which GSLs are subsequently transported to the seeds, where the 2-oxoglutarate-dependent dioxygenase ALKENYL HYDROXALKYL PRODUCING 3 (AOP3) and the SERINE CARBOXYPEPTIDASE LIKE 17 (SCPL17) can modify their structure through hydroxylation and acylation on the side chain (Xu et al. 2023; Sanden et al. 2024).

Despite the well-documented effects of ET on seed yield, oil content and storage protein composition (Angadi et al. 2000; Rashid et al. 2018; Lohani et al. 2022; Mácová et al. 2022; Devi et al. 2023; Nadakuduti et al. 2023), remarkably little is known about how seeds reshape their specialized metabolome in response to ET and how it could contribute to seed thermal-adaptation. A direct link between GSL metabolism and thermotolerance was first suggested by Ludwig-Müller et al. (2000), who showed that a glucosinolate-deficient Arabidopsis mutant exhibited impaired thermotolerance and reduced HSP90 expression after heat stress. However, the specific metabolite modifications underlying this link remain uncharacterized. Elucidating how ET modulates SM diversity and accumulation in seeds is therefore of paramount importance in the context of global warming and the need for sustainable Brassicaceae crop production.

In this study, we used multi-omic (untargeted metabolomics and transcriptomics), physiological and reverse genetic approaches to characterize the response and adaptation of *A. thaliana* seed metabolome to ET. Our findings revealed a pronounced perturbation of the flavonoid, cinnamic acid and GSL pathways under ET. In particular, we identified several novel GSLs that were sinapoylated and benzoylated on their thioglucose moiety. The accumulation of these GSLs was particularly induced by ET together with AOP3 gene expression, suggesting that AOP3 acts as a metabolic gatekeeper for short-chain hydroxyalkyl GSL production. The characterization of aop3 mutants further demonstrated the role of AOP3 in the production of GSLs acylated on the side chain under ET. Using a reverse genetic approach, we then showed that SCPL17 and BZO1 genes are required for the synthesis of thioglucose sinapoylated and/or benzoylated GSLs. Finally, we observed that the accumulation of acylated GSLs in seeds of 85 *A. thaliana* accessions corelates with the mean annual temperature of their origin sites. Taken together these results demonstrate that thioglucose GSL acylation has an important role in *A. thaliana* seed metabolic responses and adaptation to ET.

## RESULTS

### The specialized metabolite landscape is shaped by elevated temperature during A. thaliana seed development

Untargeted metabolomic (LC-MS/MS) analysis of polar and semi-polar SMs was carried out on *A. thaliana* Columbia-0 (Col-0) seeds harvested at six developmental stages (globular, transition, torpedo, bent cotyledon, mature-green and dry seed) from plant grown either under control (16h of light – 21°C/ 8h of dark – 19°C) or ET (16h of light – 27°C/ 8h of dark – 24°C) condition (Barreda et al. 2025a). The data obtained revealed how the SM landscape was affected by ET (Barreda et al. 2025a). Several metabolic categories were strongly modulated by ET during seed development (**Figure 1a; Supplemental Dataset 1a**). The highest inductions were observed for glutathione derivatives (3 SMs; log_2_ [ET/Ctr] = 4.99), carbohydrates (34 SMs; log_2_ [ET/Ctr] = 3.81), choline derivatives (3 SMs; log_2_ [ET/Ctr] = 3.49) and GSLs (34 SMs; log_2_ [ET/Ctr] = 2.70) metabolic categories (**Figure 1a; Supplemental Dataset 1a**). Overall, in this data paper (Barreda et al. 2025a) we had previously showed that 73.2 % (1274 differentially accumulated metabolites [DAMs]) and 30.9 % (537 DAMs) of the SMs were affected by stage and growing condition, respectively, revealing a strong effect of ET on the seed specialized metabolome (**Figure 1b**). Next, the DAMs induced (log_2_ [ET/Ctr] ≥ 1) or repressed (log_2_ [ET/Ctr] ≤ −1) by ET were identified for each seed developmental stage among the 537 DAMs affected by the growing condition (**Figure 1c; Supplemental Datasets 1b and 1c**). Interestingly, most DAMs were induced rather than repressed by ET during seed development (**Figure 1c**). At the bent cotyledon stage, ET strongly induced the accumulation of flavonoids, cinnamic acids and GSLs with 46.5%, 23.4% and 29.4% of compounds in each class, respectively, showing increased levels. (**Figure 1d**). Furthermore, GSLs (e.g. 3-methylsulfinylpropyl GSL [3MSOP GSL; Glucoiberin], 4-methylsulfinylbutyl GSL [4MSOB GSL; Glucoraphanin] and 3-benzoyloxypropyl GSL [3BZO GSL; Glucomalcomiin]), cinnamic acids (Sinapoylcholine 4-O-hexoside and Hydroxyferuloylcholine) and flavonoids (e.g. Isorhamnetin-3,7-dirhamnoside and Procyanidin B2) were the most represented specialized metabolic categories among the top-15 ET-induced SMs associated to a metabolic category (**Supplemental Dataset 1d**).

**Figure 1:**
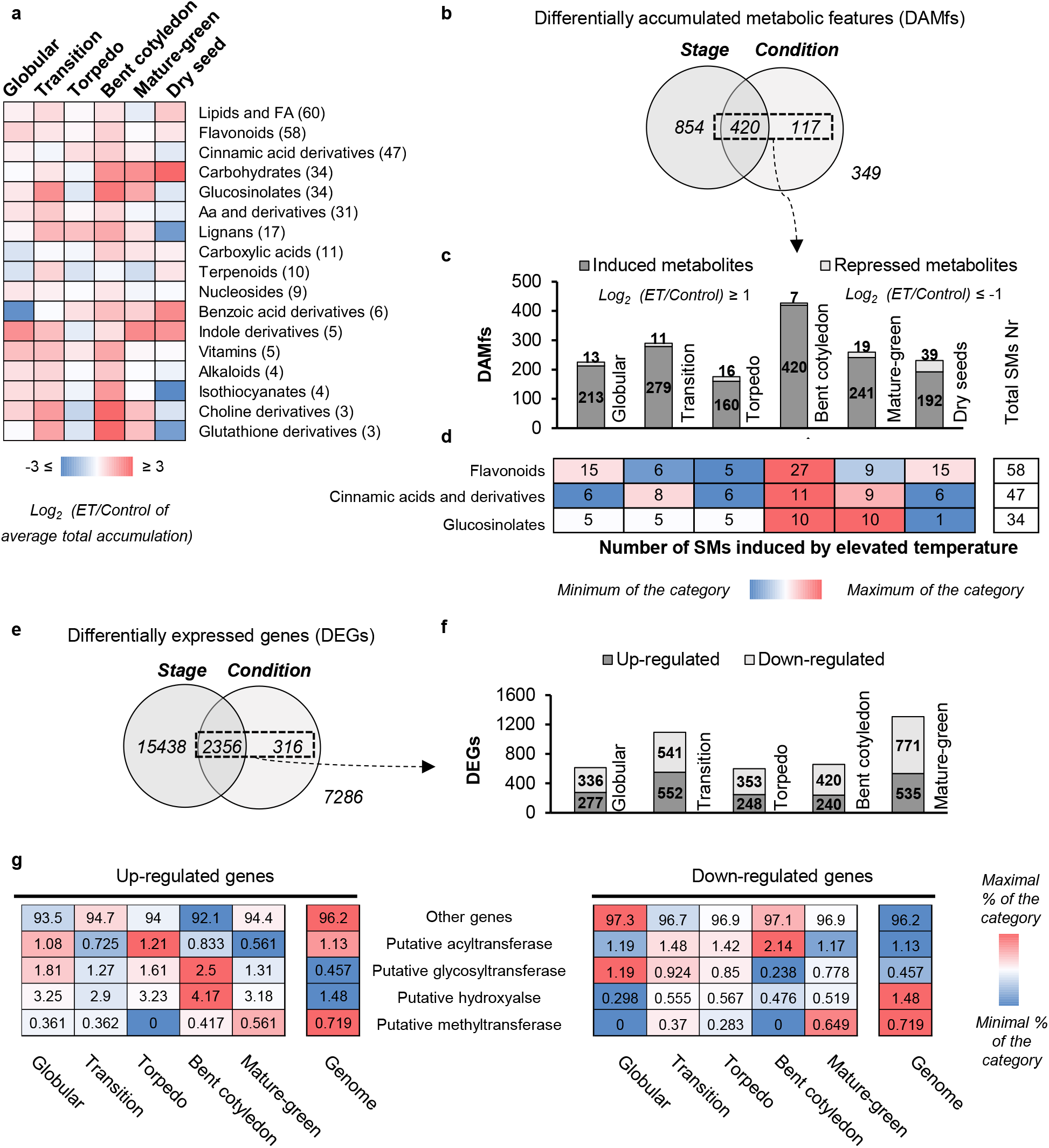
Specialized metabolites and genes affected by elevated temperature during *Arabidopsis thaliana* seed development. a) Log_2_([elevated temperature/control] average total accumulation) of metabolic categories (n≥3) at six different seed developmental stages. b) Untargeted metabolomic data statistical analysis. c) Differentially accumulated metabolic features during seed development. d) Number of major metabolic categories (Flavonoids, Cinnamic acids and derivatives, and Glucosinolates) induced by elevated temperature at each seed developmental stage. e) Transcriptomic data statistical analysis. f) Differentially expressed genes during seed development. g) Percentages of genes coding for enzymes putatively involved in specialized metabolite modifications (Acyltransferases, Glycosyltransferases, Hydroxylases and Methyltransferases) induced, and repressed, at each seed developmental stage by elevated temperature.

### The seed transcriptome is strongly impacted by elevated temperature during *A. thaliana* seed development

In parallel, RNAseq data obtained from the series of developing seeds produced under control and ET conditions were used to evaluate the effect of ET on *A. thaliana* Col-0 seed transcriptome (Barreda et al. 2025a). In total, 70 % (17794 differentially expressed genes [DEGs]) and 10.5 % (2672 DEGs) of the genes had previously been reported to be affected by stage and condition respectively (Barreda et al. 2025a) (**Figure 1e**). The genes induced (log_2_ [ET/Ctr] ≥ 1) and repressed (log_2_ [ET/Ctr] ≤ −1) by ET were identified for each developmental stage among the 2672 DEGs affected by the growing conditions (**Figure 1e; Supplemental Dataset 2a**). Unlike the metabolomic response, the number of genes up- or down-regulated by ET was similar (**Figure 1f**). Moreover, the highest number of ET-modulated DEGs were observed at transition (552 up-regulated and 541 down-regulated) and mature-green (535 up-regulated and 771 down-regulated) stages (**Figure 1f**). GO term enrichment analyses revealed that ET-induced DEGs were associated with stress-response pathways (e.g. « response to heat », « protein folding », « response to hydrogen peroxide », or « heat acclimation » GOs), confirming the strong impact of ET on the seed transcriptome (**Supplemental Figure S1** and **Supplemental Dataset 2b**). Likewise, several genes coding for heat shock proteins (e.g. *HSP23*.*6, HSP23*.*5, HSP70*.*13, HSP17*.*4* and *HSP17*.*6*) were identified among the most ET-induced genes during seed development, together with the *METHIONINE SULFOXIDE REDUCTASE B9 (MSRB9)* gene coding for an enzyme involved in the reduction of sulfoxide methionine related to oxidative stress (**Supplemental Dataset 2b**) (Li et al. 2012; Rey and Tarrago 2018). Next, genes involved in major A. thaliana seed specialized metabolome pathways, notably flavonoids, GSLs, and cinnamic acids, were identified among the 2672 DEGs affected by ET. Surprisingly, few genes involved in the biosynthesis, transport, and regulation (including transcription factors) of flavonoids (11 DEGs), GSLs (8 DEGs), and cinnamic acids (4 DEGs) showed significant modulation in their expression under ET (**Supplemental Figure S2**). The majority of those DEGs were down-regulated by ET, except for *MYB111* and *PRODUCTION OF ANTHOCYANIN PIGMENT 1 (PAP1)* (coding for transcription factors controlling flavonol biosynthesis), two genes involved in GSL side-chain modifications (*CYP81F3 MONOOXYGENASE* and *AOP3*), and one gene involved in GSL degradation (*PYK10*) (**Supplemental Figure S2**).

It has recently been shown that the type of modification (e.g. hydroxylation or acylation) occurring on some SM core structure correlates with the environmental conditions (i.e. stresses) to which seeds of the Brassicaceae species *C. sativa* are subjected (Boutet et al. 2022). These findings suggested that SM modifications may contribute to plant and seed stress responses and adaptation. To explore this hypothesis, we considered candidate genes coding for enzymes putatively involved in SM modifications, i.e. hydroxylases, glycosylases, methyltransferases or acyltransferases (as previously reported (Barreda et al. 2024)), among the 2672 DEGs affected by ET (**Supplemental Dataset 2c**). Following, the percentages of genes coding for enzymes of each of those categories, which were either up-regulated or down-regulated by ET, were calculated for each seed developmental stage and compared to their percentage in the genome (**Figure 1g**). Interestingly, under ET, genes coding for both putative SM hydroxylases and glycosyltransferases were globally induced by ET (**Figure 1g**).

### Multi-omics suggest that *AOP3* pathway and associated glucosinolates play roles in seed response to elevated temperature

To identify SM modification pathways involved in *A. thaliana* seed response to ET, we used our previously published multi-omic integrative analysis (Barreda et al. 2025a). *ALKENYL HYDROXALKYL PRODUCING 3 (AOP3)* gene expression was highly induced by ET from torpedo to mature-green stage and correlated with the accumulation of SMs belonging to several metabolic categories, including many GSLs (**Figure 2**). Previous studies showed that AOP3 codes for a 2-oxoglutarate dependent dioxygenase involved in (short) methylsulfinylalkyl-GSL side chain hydroxylation, resulting in the formation of hydroxyalkyl GSLs (Kliebenstein et al. 2001). Consistently, our analysis highlighted that the accumulation of a putative 3BZO GSL (Glucomalcomiin), which is a derivative of the hydroxyalkyl GSL 3-hydroxypropyl GSL (3OHP GSL), was highly correlated with *AOP3* expression (**Figure 2**). In addition, *AOP3* expression was correlated with 8 other GSLs including the 3 annotated as follows: glucolesquerellin (n333), methylsulfanylheptyl GSL (n99) and 5-methylhexyl GSL (n708) (**Figure 2**). For the 5-remaining unannotated GSLs, we further analysed their MS^2^ spectra and determined their putative structure. The GSL with *m/z* 700.1384 was annotated as 6’-sinapoyl-4-benzoyloxybutyl GSL (6’-sin-4BZO GSL) and the GSL with *m/z* 686.1227 as 6’-sinapoyl-3-benzoyloxypropyl GSL (6’-sin-3BZO GSL), which correspond to new side-chain benzoylated hydroxyalkyl GSLs that are sinapoylated on their thioglucose moiety (**Figure 2**). 6’-sinapoyl-GSLs present MS^2^ diagnostic ions indicating the presence of a sinapoyl group on the thioglucose (*m/z* 447 [C17H19O12S^−^], 465 [C17H21013S^−^], 481 [C17H21O12S2^−^])(Shi et al. 2017) that are present in the MS^2^ of both 6’-sin-4BZO GSL and 6’-sin-3BZO GSL presented in this study (**Figure 2**). Furthermore, putative 6’-benzoyl-4-methylthiobutyl GSL (6’-bnz-4MTB GSL; m/z 598.1065), which corresponds to the thioglucose benzoylated form of 4MTB GSL, that is situated upstream the 4MSOB GSL precursor of AOP3, was also annotated among the GSLs correlating with *AOP3* expression (**Figure 2**). The induction of *AOP3* expression and accumulation of related newly identified acylated GSLs by ET suggested a strong involvement of this pathway in A. thaliana Col-0 seed response to ET, which was further investigated.

**Figure 2:**
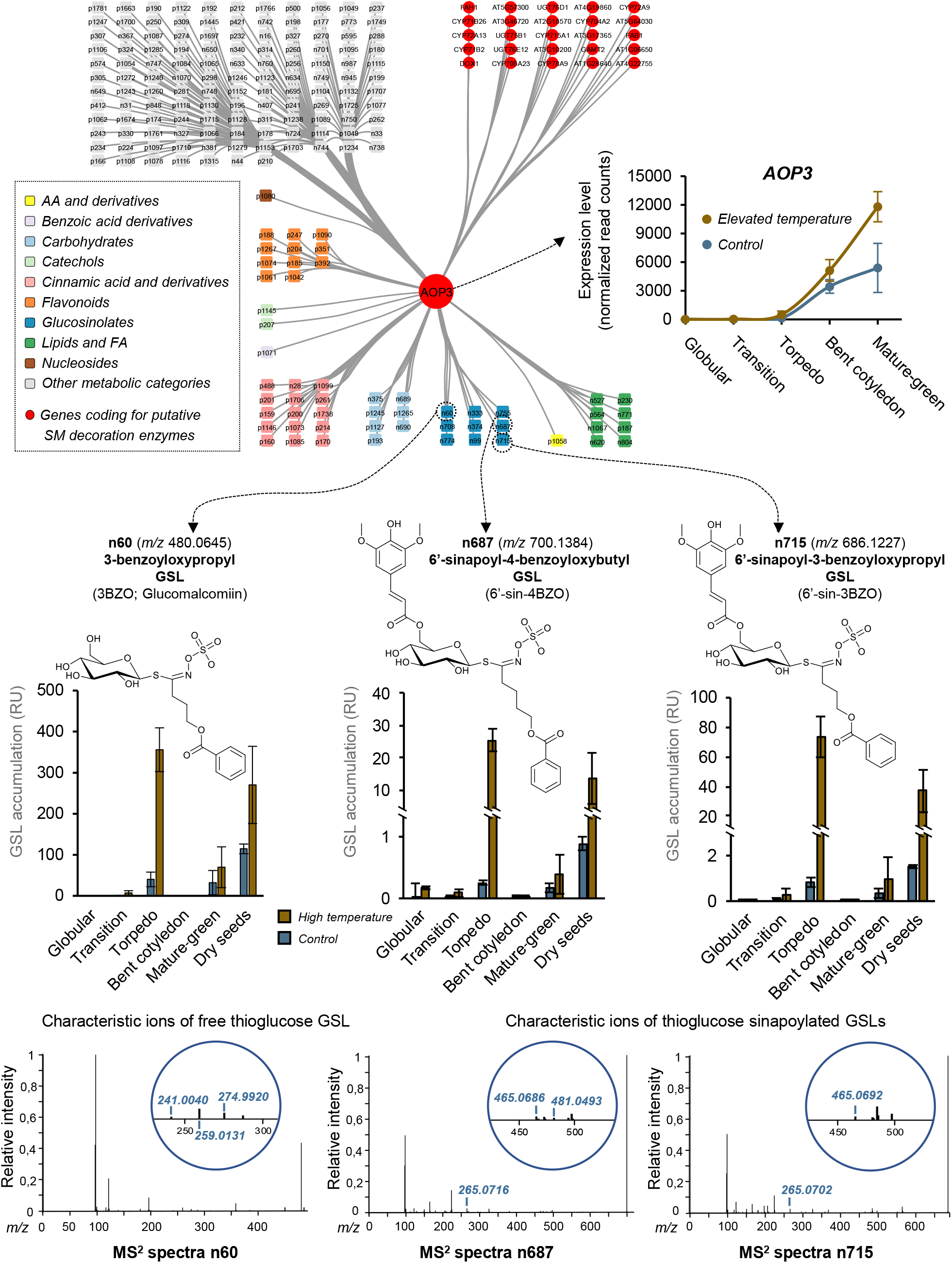
Correlation of *AOP3* gene expression with the accumulation of thioglucose sinapoylated glucosinolates. The accumulations of several specialized metabolites were found to correlate with *AOP3* expression during seed development. In particular, *AOP3* expression and several glucosinolates accumulations were found to be induced by elevated temperature during seed development.

### Comparative analysis of *aop3* and wild-type genotypes demonstrate the presence of novel thioglucose-acylated glucosinolates accumulating in developing seeds under elevated temperature conditions

Untargeted metabolomic (LC-MS/MS) analysis of polar/semi-polar SMs was performed on developing seeds (torpedo, mature-green, and dry seed stages) of *A. thaliana* aop3-1 knock-out (SALK_001655C) and wild-type lines grown under control or ET condition. A total of 1440 peaks of putative SMs were detected and a molecular networking approach was used to cluster them based on MS^2^ spectra, improving the annotation of unknown metabolites (**Supplemental Datasets 3a and 3b**)(Olivon et al. 2018; Boutet et al. 2022; Barreda et al. 2025a). Principal component analysis (PCA) conducted on untargeted metabolomic data allowed to separate the samples according to the genotype and the growing condition of seed development, however seed stage was the most discriminating factor of the samples (**Supplemental Figure S3**). Pairwise analyses (log_2_ [ET/Ctr] ≥ |1|; pvalue ≤ 0.05) were conducted to identify SM categories induced or repressed by ET in *aop3-1* and the wild type, at different seed developmental stages (torpedo, mature-green and dry seed) (**Supplemental Figure S4**). Among all metabolic categories, GSLs showed the highest induction by ET, with 3 to 20 GSLs induced by ET depending on the stage and genotype considered, confirming their induction by ET response. In addition, cinnamic acids (from 4 to 9), flavonoids (from 4 to 11) and carbohydrates (from 3 to 15) were induced by ET (**Supplemental Figure S4**). Interestingly, solely 3 GSLs were induced in the wild type at torpedo stage whereas 17 GSLs were induced at this stage in *aop3-1*, suggesting that the difference of induction of GSL accumulation in these two genotypes may start quite early during seed development. Mature-green was the stage at which the higher number of GSLs were induced by ET with 17 and 20 GSLs induced in the wild type and *aop3-1*, respectively (**Supplemental Figure S4**). At mature-green stage, the majority of ET-induced GSLs with putative annotation were newly described methylthiosulfinyl and methylthiobutyl GSLs acylated, harbouring sinapoyl [sin] or benzoyl [bnz], on their thioglucose (6’-acyl-GSL). The 6’-sinapoyl-GSLs described here presented at least several MS^2^ diagnostic ions indicating the presence of a sinapoyl group on the thioglucose described previously (Shi et al. 2017) whereas the 6’-benzoyl-GSLs presented at least several MS^2^ diagnostic ions indicating the presence of a benzoyl group on the thioglucose (*m/z* 345 [C13H13O9S^−^], 363 [C13H15010S^−^], 379 [C13H15O9S2^−^]).

Surprisingly, aop3-1 still accumulated several thioglucose-acylated GSLs bearing mid-long methylthioalkyl or methylsulfinyl side chains, substrates that are not processed by AOP3. This demonstrates that SCPL17-mediated thioglucose acylation can occur independently of AOP3-mediated side-chain hydroxylation and operates on a broader range of GSL substrates than previously recognized. Specifically, these metabolites included 6’-sin-7-methylthioheptyl [6’-sin-7MTH] GSL, 6’-sin-5-methylthiopentyl [6’-sin-5MTP] GSL, 6’-bnz-6-methylthiohexyl [6’-bnz-6MTH] GSL, 6’-bnz-7-methylthioheptyl [6’-bnz-7MTH] GSL, 6’-bnz-8-methylthiooctyl [6’bnz-8MTO] GSL, 6’-sin-8-methylsulfinyloctyl [6’-sin-8MSOO] GSL, 6’-bnz-8-methylsulfinyloctyl [6’-bnz-8MSOO] GSL and 6’-sin-6-methylthiohexyl [6’-sin-6MTH] GSL (**Figure 3**). Similarly, some 6’-acyl-GSLs such as 6’-sin-5MTP GSL, 6’-bnz-6MTH GSL, 6’-sin-7MTH GSL, 6’-sin-4MTB GSL and 6’-sin-4BZO GSL were found to be induced in aop3-1 and/or the wild type at dry seed stage (**Supplemental Figure S5**). Moreover, the GSLs acylated on their side-chain, such as 6’-sin-4BZO, were much less accumulated in the aop3-1 mutant (**Supplemental Figure S5**). In addition, all those 6’-acyl-GSLs are specifically accumulated in mature-green and dry seeds, while they were not detected in leaves (**Supplemental Figure S6**).

**Figure 3:**
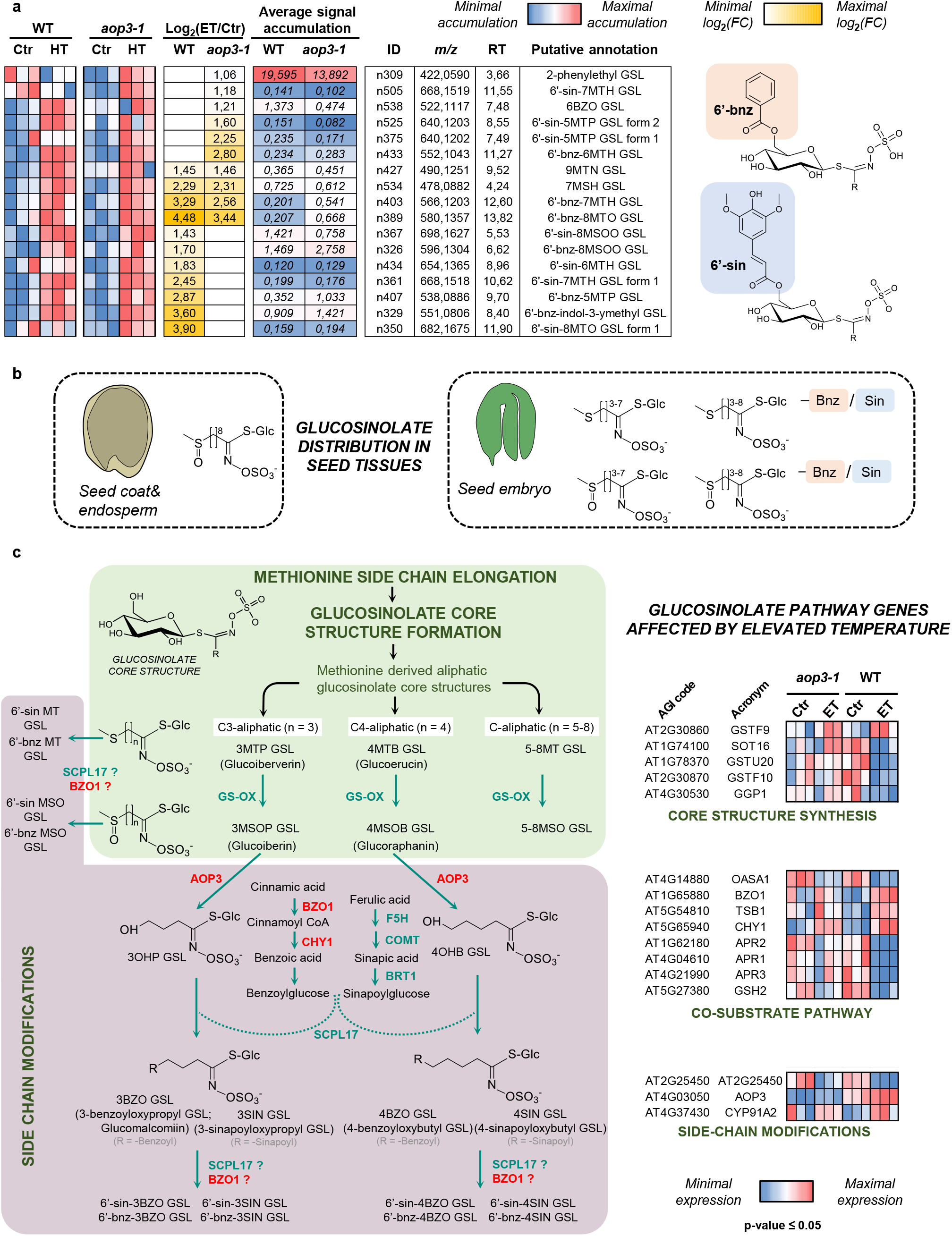
Glucosinolates and related biosynthetic genes are affected by elevated temperature in *aop3-1* mutant and wild-type at mature-green stage of seed development. a) Relative accumulation heatmaps, log_2_([elevated temperature/control] average total accumulation) (log_2_[ET/Ctr]), control accumulation averages, metabolite identity (ID), mass/charge ratio (*m/z*), retention time (RT), putative annotation and spectral annotation are listed for each glucosinolate. b) Distribution of glucosinolates and related degradation products between Seed coat & endosperm (SC) and seed embryo (SE) in *aop3-1* mutant and wild-type dry seeds developed under control and elevated temperature. Abbreviations: Sin, sinapoyl; Bnz, benzoyl. c) Methionine derived aliphatic glucosinolate biosynthesis pathway highlighting the genes affected by elevated temperature in *aop3-1* and wild type at mature-green seed developmental stage. Hydroxylases, methyltransferases and acyltransferases are represented in orange, green and blue respectively. Abbreviations of glucosinolate (GSL) names: 3MTP GSL, 3-methylthiopropyl GSL (Glucoiberverin); 4MTB GSL, 4-methylthiobutyl GSL (Glucoerucin); 5MTP GSL, 5-methylthiopentyl GSL (Glucoberteroin); 6MTH GSL, 6-methylthiohexyl GSL (Glucolesquerellin); 7MTH GSL, 7-methylthioheptyl GSL; 8MTO GSL, 8-methylthiooctyl GSL; 3MSOP GSL, 3-methylsulfinylpropyl GSL (Glucoiberin); 4MSOB GSL, 4-methylsulfinylbutyl GSL (Glucoraphanin); 5MSOP GSL, 5-methylsulfinylpentyl GSL (Glucoalyssin); 6MSOH GSL, 6-methylsulfinylhexyl GSL (Glucohesperin); 7MSOH GSL, 7-methylsulfinylheptyl GSL (Glucoibarin); 8MSOO GSL, 8-methylsulfinyloctyl GSL (Glucohirsutin); 3OHP GSL, 3-hydroxypropyl GSL; 4OHB GSL, 4-hydroxybutyl GSL; 3BZO GSL, 3-benzoyloxypropyl GSL (Glucomalcomiin); 3SIN GSL, 3-sinapoyloxypropyl GSL; 4BZO GSL, 4-benzoyloxybutyl GSL; 4SIN GSL, 4-sinapoyloxybutyl GSL; 6’-bnz-3BZO GSL, 6’-benzoyl-3-benzoyloxypropyl GSL; 6’-sin-3BZO GSL, 6’-sinapoyl-3-benzoyloxypropyl GSL; 6’-bnz-3SIN GSL, 6’-benzoyl-3-sinapoyloxypropyl GSL; 6’-sin-3SIN GSL, 6’-sinapoyl-3-sinapoyloxyproyl GSL; 6’-bnz-4BZO GSL, 6’-benzoyl-4-benzoyloxybutyl GSL; 6’-sin-4BZO GSL, 6’-sinapoyl-4-benzoyloxybutyl GSL; 6’-bnz-4SIN GSL, 6’-benzoyl-4-sinapoyloxybutyl GSL; 6’-sin-4SIN GSL, 6’-sinapoyl-4-sinapoyloxybutyl GSL; 6’-sin GSL, 6-’thioglucose-sinapoyl GSL; 6’-bnz GSL, 6’-thioglucose-benzoyl GSL. Gene coding enzymes names : GS-OX, flavin-monooxygenase glucosinolate oxygenase; AOP3, alkenyl hydroxalkyl-producing 3; CHY1, cinnamoyl-CoA hydrolase; SCPL17, serine carboxypeptidase-like 17; BZO1, benzoyloxy GSL 1; F5H, ferulic acid 5-hydroxylase; COMT, caffeate/5-hydroxyferulate 3-O-methyltransferase; BRT1, bright trichomes 1; CYP, cytochrome P450; GST, glutathione S-transferase; SOT, sulfotransferase; GGP1, γ-glutamyl peptidase 1; OASA1, cysteine synthase 1; TSB1, tryptophan synthase β chain 1; APR, 5′-adenylylsulfate reductase 1; GSH, glutathione synthase.

To evaluate the potential effect of metabolite acylation and/or ET on GSL localisation, seed coat plus endosperm (SC) and embryo tissues (SE) were dissected of wild-type and *aop3-1* dry seeds produced under control or ET condition and subsequently profiled by untargeted metabolomics conditions (**Figure 3b; Supplemental Figure S7; Supplemental Dataset 4**). GSLs and nitrile degradation products were mainly accumulated in the SE, while isothiocyanates (ITCs) were accumulated in the SC. The non-acylated long-chain GSLs (C≥8; OH-8MSOO GSL and 8MSOO GSL) were accumulated in the SC while their sinapoylated and benzoylated forms were accumulated in the embryo (6’-sin-8MSOO GSL, 6’-bnz-8MSOO GSL, 6’-bnz-8MTO GSL and 6’-sin-8MTO GSL) (**Figure 3b**). In a complementary experiment including a second T-DNA insertion mutant line of *AOP3* (SALK_022752C, referred to as *aop3-2*) and wild-type genotypes, untargeted metabolomic performed on dry seeds confirmed that several of the GSLs induced by ET in the wild type and in *aop3-2* were acylated on their thioglucose moiety (**Supplemental Figures S8 and S9; Supplemental Datasets 5a and 5b**).

### Transcriptomic remodelling induced by elevated temperature is impaired in *aop3* seeds

RNAseq analyses were conducted with *aop3-1* and wild-type seeds (mature-green stage) produced either under control or ET conditions (**Supplemental Dataset 6a**). The PCA performed allowed to separate most of the samples according to the genotype and condition (**Supplemental Figure S10a**). The PC1 axis, explaining 22.2% of the variability, allowed to separate the samples according to the condition. Interestingly, the transcriptome of *aop3-1* seeds was less affected by ET compared to that of the wild type. Wild-type seeds more than tripled the number of ET-modulated DEGs compared to the *aop3-1* mutant. In particular, ET down-regulated 3657 and 1269 genes and up-regulated 3183 and 779 genes in wild-type and *aop3-1* seeds, respectively (**Supplemental Figure S10b; Supplemental Dataset 6b**). Genotype-specific and common genes were identified for both ET up-regulated and down-regulated genes (**Supplemental Figure S10c and Supplemental Dataset 6c**).

The effect of ET was then evaluated on the expression of GSL-related genes (**Supplemental Figures S11a and S11b**). Genes involved in GSL degradation and transport were affected by ET. In particular, among the genes coding for myrosinases, that are involved in the removal of GSL thioglucose moiety, *TGG1* (*THIOGLUCOSEHYDROLASE 1; BGLU38*) was up-regulated by ET in the wild type whereas *PEN2 (PENETRATION 2; BGLU26), BGLU28 (BETA GLUCOSIDASE 28)* and *DIN2 (DARK INDUCIBLE 2; BGLU30)* were down-regulated (**Supplemental Figure S11a**). The epithiospecificer *ESM1 (EPITHIOSPECIFIER MODIFIER 1)*, involved in the formation of epithionitriles, was down-regulated in the wild type upon ET. Finally, the majority of nitrilases and nitrile specifier proteins (*NITRILASE 1 [NIT1], NIT2, NIT3, NSP1* and *NITRILE SPECIFIER 3*) were down-regulated by ET whereas *NIT4* and *NSP2* were up-regulated by ET (**Supplemental Figure S11a**). Furthermore, genes coding for the transporters involved in GSL seed loading were differentially expressed upon ET (Sanden et al. 2024). *GTR2 (GLUCOSINOLATE TRANSPORTER 2)* was up-regulated by ET in both genotypes, while *UMAMIT30 (USUALLY MULTIPLE ACIDS MOVE IN AND OUT TRANSPORTERS 30)* was induced by ET in the wild-type (**Supplemental Figure S11a**). Inversely, *UMAMIT29* was down-regulated by ET in both genotypes while *GTR3* was down-regulated in the wild type (**Supplemental Figure S11b**).

The transcriptomic analysis confirmed that *AOP3* was induced by ET in the wild type (log_2_[ET/Ctr]_wild type_=0.65) (**Supplemental Dataset 5b**) whereas its expression was significantly reduced in the *aop3-1* mutant in both ET (log_2_[wild type/*aop3-1*]_ET_=1.91) and control condition (log_2_[wild type/*aop3-1*]_Ctr_=1.58) conditions (**Figure 3c; Supplemental Dataset 6d**). Furthermore, while several genes involved in co-substrate pathways (e.g. *CYSTEIN SYNTHASE 1 [OASA1], APS REDUCTASE 1 [APR1], APR2, APR3* and *GLUTATHIONE SYNTHETASE 2 [GSH2]*) were repressed by ET in wild-type seeds, the genes *TRYPTOPHAN SYNTHASE BETA-SUBUNIT 1 (TSB1), BENZOYLOXYGLUCOSINOLATE 1 (BZO1)* and *3-HYDROXYISOBUTYRYL-COA HYDROLASE 1 (CHY1)* were induced by ET in the wild type (**Figure 3c**). BZO1 and CHY1 are involved in the production of the benzoyl donor that is used by SCPL17 (SERINE CARBOXYPEPTIDASE LIKE 17) acyltransferase to add benzoyl groups to the side-chains (Kliebenstein et al. 2007; Ibdah and Pichersky 2009; Lee et al. 2012). Those results could therefore suggest that the higher accumulation of 6’-benzoyl-GSLs may be linked to the induction of *CHY1* and *BZO1* expression. SCPL17 is also involved in the transfer of sinapoyl groups to GSL side-chains (Lee et al. 2012). Furthermore, consistently with the higher accumulation of 6’acyl-GSLs in seeds, *SCPL17, BZO1* and *AOP3* are specifically expressed in *A. thaliana* seeds (**Supplemental Figure S12**). Hence, *SCPL1*7 appeared as a good candidate for the sinapoylation and benzoylation of GSL thioglucose moieties.

### SCPL17 and BZO1 are involved in thioglucose acylation of glucosinolates

To further study the genetic regulation and involvement in ET response of GSL side-chain and thioglucose sinapoylation and benzoylation, knock-out mutants of *SCPL17* (*scpl17-1* and *scpl17-2*), *BZO1* (*bzo1-4* and *bzo1-6*), *AOP3* (*aop3-1*) and wild-type lines were grown in control and ET conditions. Dry plant weight and seed number per silique were assessed in wild-type and mutant lines. Under ET condition, wild-type, *aop3-1* and *scpl17-1* exhibited increased dry plant weight compared to control condition (Mann-Whitney tests; p-value ≤ 0.05) (**Supplemental Figure S13a**). Furthermore, seed number per silique was reduced under ET compared to control condition in *aop3-1, bzo1-4, bzo1-6* and *scpl17-1* mutants, but not in the wild-type (Mann-Whitney tests; p-value ≤ 0.05) (**Supplemental Figure S13b**). The total fatty acid content of dry seeds was also analysed. As highlighted by an increased C18:2/C18:3 ratio measured under ET, the production of triunsaturated fatty acids was decreased by ET (**Supplemental Figure S13c, Supplemental Dataset 7**). Nevertheless, within each growing condition, fatty acid composition did not differ significantly between genotypes. Besides, DPPH (2,2-diphenyl-1-picrylhydrazyl) test was performed to evaluate the antioxidant capacities of seed SMs extracts of the different genotypes and temperature conditions (**Supplemental Figure S13d**). EC50 values (effective concentration which scavenges 50% of DPPH radical) were generally higher for the SM extracts of ET developed seeds (compared to control developed seeds), indicating that their antioxidant capacities were lower (**Supplemental Figure S13d**). However, there were no EC50 values differences among the genotypes (**Supplemental Figure S13d**).

To evaluate the impact of the genotype and/or the ET stress condition on seed metabolite composition, untargeted metabolomic analysis was performed on *scpl17, bzo1, aop3* and wild-type lines. A total of 779 SM ions were detected and annotated (**Supplemental Datasets 8a and 8b**) with PCA of the untargeted metabolomic data revealing two genotype-dependent clusters that separated *aop3* and *scpl17* from *bzo1* and the wild type (**Figure 4a**).

**Figure 4:**
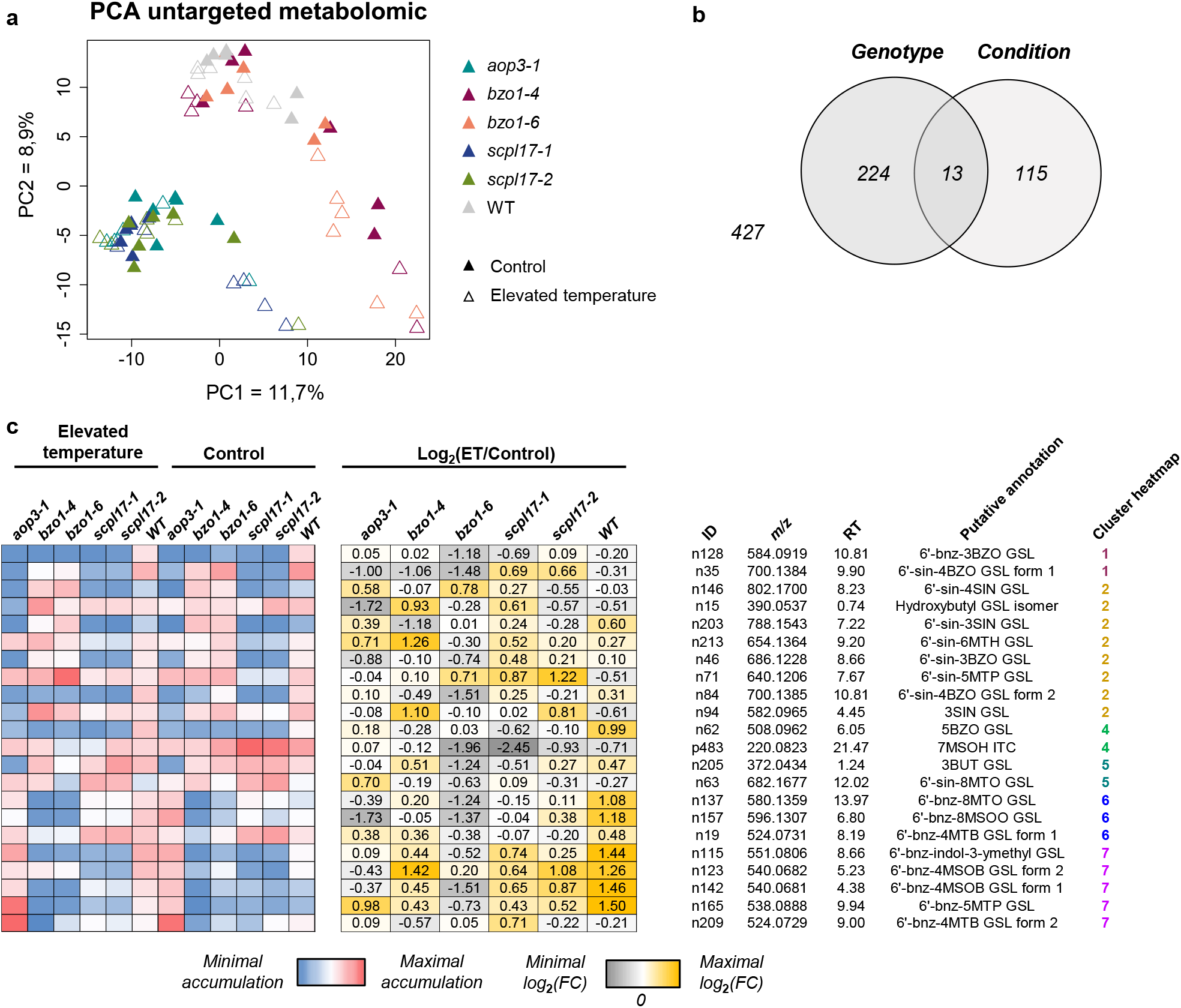
Untargeted metabolomic analysis of *aop3, bzo1* and *scpl17* mutants and wild-type dry seeds developed under elevated temperature or control condition. a) Principal component analysis on untargeted metabolomic data. b) Statistical analysis on untargeted metabolomic data. c) Glucosinolates and related degradation products affected by elevated temperature in *aop3, bzo1, scpl17* mutants and wild type at dry seed developmental stage. Relative accumulation heatmaps, log_2_(elevated temperature/control average total accumulation) (log_2_[ET/Ctr]), metabolite identity (ID), mass/charge ratio (*m/z*), retention time (RT), putative annotation and number of heatmap cluster are listed for each glucosinolate and degradation product. Abbreviations of glucosinolate (GSL) and isothiocyanate (ITC) names: 3BZO GSL, 3-benzoyloxypropyl GSL (Glucomalcomiin); 3SIN GSL, 3-sinapoyloxypropyl GSL; 4BZO GSL, 4-benzoyloxybutyl GSL; 4SIN GSL, 4-sinapoyloxybutyl GSL; 6-’bnz-3BZO GSL, 6’-benzoyl-3-benzoyloxypropyl GSL; 6’-sin-3BZO GSL, 6’-sinapoyl-3-benzoyloxypropyl GSL; 6’-sin-3SIN GSL, 6’-sinapoyl-3-sinapoyloxyproyl GSL; 6’-sin-4BZO GSL, 6’-sinapoyl-4-benzoyloxybutyl GSL; 6’-sin-4SIN GSL, 6’-sinapoyl-4-sinapoyloxybutyl GSL; 6’-sin-6MTH GSL, 6’-sinapoyl-6-methylsulfinylhexyl GSL; 6’-sin-5MTP GSL, 6’-sinapoyl-5-methylpentyl GSL; 5BZO GSL, 5-benzoyloxy GSL; 7MSOH GSL, 7-methylsufinylheptyl GSL (Glucoibarin); 7MSOH ITC, 7-methylsufinylheptyl ITC; 3BUT GSL, 3-butenyl GSL; 6’-sin-8MTO GSL, 6’-sinapoyl-8-methylthiooctyl GSL; 6’-bnz-8MTO GSL, 6’-benzoyl-8-methylthiooctyl GSL; 6’-bnz-8MSOO GSL, 6’-benzoyl-8-methylsulfinyloctyl GSL; 6’-bnz-4MTB GSL, 6’-benzoyl-4-methylthiobutyl GSL; 6’-bnz-indol-3-ymethyl GSL, 6’-benzoyl-indol-3-ymethyl GSL; 6’-bnz-4MSOB GSL, 6’-benzoyl-4-methylsufinylbutyl GSL; 6’-bnz-5MTP GSL, 6’-benzoyl-5-methylpentyl GSL.

Statistical analyses allowed the identification of 224 SMs modulated by the genotype (*aop3, scpl17, bzo1*, wild type), 115 SMs affected by the condition (control, ET), and 13 SMs impacted by both factors (**Figure 4b; Supplemental Dataset 8c and 8d**). Hierarchical clustering analysis allowed to regroup the SMs into 7 distinct clusters according to their accumulation patterns (**Supplemental Figure S14**). Cinnamic acids (39 DAMs), and GSLs, along with their related degradation products (37 DAMs), were the metabolic categories that exhibited the most pronounced differences in accumulation among the wild type and mutants (**Figure 4c; Supplemental Figure S14**).

In particular, GSLs acylated on both their thioglucose moiety and side-chain (e.g. 6’-bnz-3BZO GSL, 6’-sin-4BZO GSL, 6’-sin-4SIN GSL, 6’-sin-3SIN GSL, 6’-sin-3BZO GSL), and the corresponding free-thioglucose forms (**Figure 4c**, clusters 1 and 2), showed lower accumulation in *aop3-1, scpl17-1* and *scpl17-2* compared to the wild type, confirming the role of AOP3 in the hydroxylation and of SCPL17 in the subsequent acylation of short aliphatic GSLs (C3 and C4) side-chains. Moreover, the 6’-sinapoyl-methylthioalkyl GSLs, 6’-sin-6MTH GSL and 6’-sin-5MTP GSL, accumulated to lower levels in the seeds of both *scpl17* mutant lines, while they showed similar accumulation level between *aop3-1* and the wild type. This suggested that SCPL17 is involved in the sinapoylation of GSL thioglucose moieties. Interestingly, the majority of 6’-sin-GSLs of clusters 1 and 2 (**Figure 4c**), were more accumulated in *bzo1* than in the wild type, suggesting that 6’sinapoylation rate of GSLs increased when less benzoyl sources were available (**Supplemental Figure S15**). The isothiocyanates (ITCs), GSL degradation products, clustered together (cluster 4) and were characterized by a strong repression under ET (−4.16 ≥ log_2_ [ET/Ctr] ≤ −1.46) (**Figure 4c**). Several not annotated GSLs (Figure 4, cluster 5) showed lower accumulation in aop3-1, suggesting that they may be substrates of the corresponding enzyme (**Figure 4c**). Finally, the 6’benzoyl-methylthioalkyl and 6’benzoyl-methylsulfinyl GSLs (Figure 4; clusters 6 and 7; 6’-bnz-8MTO GSL, 6’-bnz-8MSOO GSL, 6’-bnz-4MTB GSL, 6’-bnz-5MTP GSL and 6’-bnz-4MTB GSL) and 6’-bnz-indol-3-ymethyl GSL were much less accumulated in *bzo1* and *scpl17* than in *aop3-1* and the wild type. Differently, 6’-bnz-4MSOB GSL accumulation was similar in *scpl17* and *aop3-1* (**Figure 4c**). Furthermore, these GSLs showed the highest induction by ET in the wild type, further emphasizing the pronounced effect of ET on the benzoylation of GSL thioglucose moieties (**Figure 4c**).

### Annual mean temperature adaptation drives variability in seed thioglucose-acylated GSL accumulation

After documenting the effect of ambient temperature responses on GSL production in the seeds of a given accession (Col-0) grown under different temperature conditions, we wanted to test whether this same parameter could be a factor that led to a lasting and constitutive adaptation of GSL production in the seeds of accessions from different geographical origins. Seed GSLs profiles of 85 accessions grown simultaneously in a greenhouse were profiled to evaluate their potential involvement in ET adaptation. Latitude and longitude coordinates of the accession origin locations, together with corresponding elevation, annual temperature average and annual precipitation were retrieved (see Material & Methods for further details). Pearson correlations between these environmental variables and GSL accumulation levels were then calculated (**Figure 5a; Supplemental Datasets 9a, 9b and 9c**). The correlation between GSL accumulation and temperature was higher than those observed between GSL and the other parameters considered (**Figure 5a; Supplemental Dataset 9d**). In particular, correlations of GSL accumulation with temperature ranged between 0.35 and −0.22 (Pearson correlation) according to the GSL considered, with 6’-sinapoyl-7-methylthioheptyl GSL (n597) presenting the strongest correlation with temperature (**Figure 5b; Supplemental Dataset 9d**). Next, GSLs were classified by structure (derived from methionine or others; with long or short side-chain; benzoylated or sinapoylated side-chains; or containing benzoylated or sinapoylated thioglucose moieties) and corresponding average correlation with temperature were determined for each category (**Figure 5c; Supplemental Dataset 9c**). Overall, the accumulation of thioglucose-acylated GSLs displayed the strongest correlations with temperature (**Figure 5c**).

**Figure 5:**
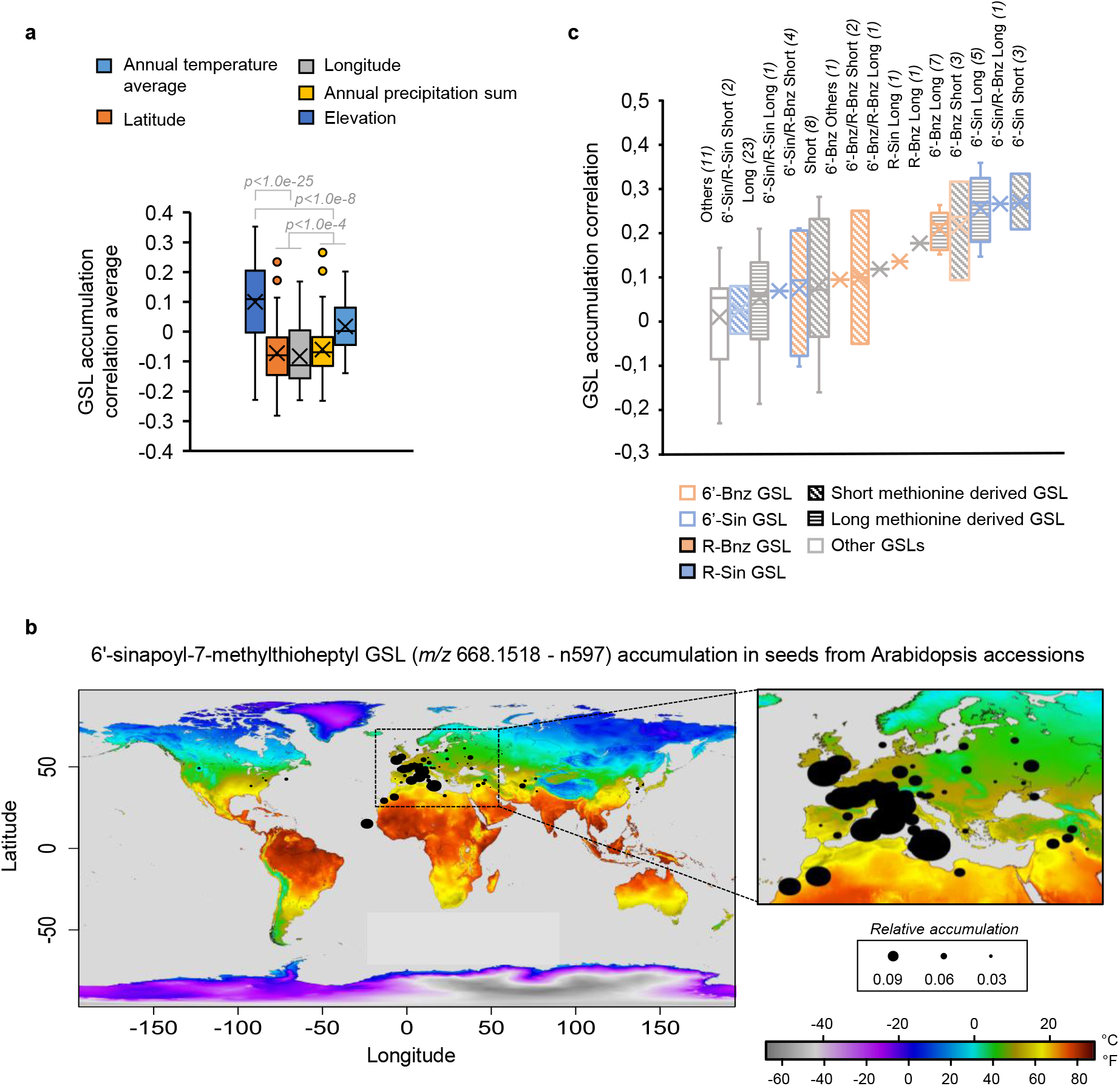
Glucosinolate accumulation in seeds from 85 *Arabidopsis thaliana* accessions. a) Average of seed glucosinolate accumulation correlation with *A. thaliana* accession origin location parameters (latitude, longitude, elevation, annual temperature average and annual precipitation sum (Kruskall-Wallis test). b) 6’-sinapoyl-7-methylthioheptyl GSL (*m/z* 668.1518 - n597) accumulation in seeds from *A. thaliana* accessions plotted on annual average temperature world map according to their origin location. The size of (black) dots is proportional to n597 accumulation. c) Correlation of average accumulation of seed glucosinolates with annual temperature of *A. thaliana* accession origins. Methionine derived GSLs benzoylated on their thioglucose (6’-Bnz GSL) and their side-chain (R-Bnz GSL) are presented with orange histogram borders and orange histogram filling respectively. Methionine derived GSLs sinapoylated on their thioglucose (6’-Sin GSL) and their side-chain (R-Sin GSL) are presented with blue histogram borders and blue histogram filling respectively. Short and long chain methionine derived GSLs are represented with diagonal and horizontal lines histogram filling respectively.

## DISCUSSION

To cope with rising temperatures associated with global warming, plants require effective strategies and mechanisms to respond and adapt to ET. SM play crucial roles in plant and seed adaptation to environmental conditions (Abbas et al. 2017; Corso et al. 2020). In addition, SM strongly influence the nutritional and physiological quality of seeds, including storage, germination, and stress resilience (Corso et al. 2021; Hanif et al. 2023). Therefore, in the context of climate change and the need for more sustainable agricultural systems, investigating the effects of ET on the seed specialized metabolome is of paramount importance. Such studies will help elucidate how seeds respond to ET and whether these metabolites contribute to seed long-term thermal adaptation.

In this study, we revealed how ET can shape the seed specialized metabolome of *A. thaliana*. During seed development, the bent cotyledon stage showed the strongest regulation of the specialized metabolome, with SMs being highly induced by ET (**Figure 1c**). Transcriptomic analysis revealed two strong gene-regulation waves at transition and mature-green stages (**Figure 1f**), indicating that both up- and down-regulation of gene expression can drive SM accumulation. Genes coding for hydroxylases and glycosylases were especially induced by ET. Although genes in flavonoid, GSL, and cinnamic acid biosynthesis were not induced, corresponding metabolites accumulated strongly, pointing to post-translational regulations. Delays between gene regulation and metabolomic changes may explain uncaptured links, highlighting the need for integrative approaches that include timing. Future work should examine transcriptomic and metabolomic responses in tissue- and cell-specific context. The apparent decoupling between gene expression and metabolite accumulation, whereby few GSL biosynthetic genes were transcriptionally induced despite strong metabolic changes, is a recurring observation in stress metabolomics (Choi et al. 2026). Bivalent histone modifications have been shown to orchestrate the temporal regulation of GSL biosynthesis during stress responses, enabling rapid metabolic shifts without sustained transcriptional activation (Choi et al. 2026). Such epigenetic mechanisms, together with post-translational regulation and metabolic channelling, may explain the discordance observed in our data and underscore the need for integrative multi-omic approaches that capture regulatory events beyond transcription.

The accumulation of indolic and aliphatic GSLs upon both high and low temperatures has been previously reported in *Brassica rapa* seedlings and in *Brassica oleracea* leaves (Valente Pereira et al. 2002; Charron and Sams 2004; Rao et al. 2021). However, little was known about how ET impacts the diversity and plasticity of GSLs in seeds. Here, we showed that GSL accumulation and acylation are strongly enhanced in *A. thaliana* seeds developed under ET (**Figure 1a and d**). While 6’sinapoyl-4-methylthiobutyl GSL (6’-sin-4MTB GSL; 6’-sin-Glucoerucin), 6’-sinapoyl-4-methylsulfinylbutyl GSL (6’-sin-4MSOB GSL; 6’-sin-Glucoraphanin) and 6’-benzoyl-4-benzoyloxybutyl GSL (6’-bnz-4BZO GSL) have been previously described in *A. thaliana* Col-0 seeds, or in cruciferous vegetable-based dietary supplements, we have identified several new thioglucose-acylated GSLs in *A. thaliana* seeds from several accessions (**Supplemental Dataset 10**) (Reichelt et al. 2002; Shi et al. 2017). These included thioglucose acylations on methylthioalkyl GSLs, methylsulfinyl GSLs but also hydroxyalkyl GSLs acylated on their side-chain or not. Previous nuclear magnetic resonance (NMR) analyses on 6’-bnz-4BZO GSL revealed that benzoyl esterification occurs on the C-6’ of the thioglucose moiety, suggesting that the other thioglucose-acylated GSLs observed in this study could be also acylated on their C-6’ (Reichelt et al. 2002). However, isomers (i.e. same *m/z* but different retention times) were detected for some of the putative thioglucose-acylated GSLs, suggesting an acylation on alternative C-thioglucose positions.

The accumulation of these newly identified thioglucose-acylated GSLs was linked to both responses and adaptation to ET. Their levels increased significantly under ET, and their abundance in seeds from natural *A. thaliana* populations adapted to specific environments showed a strong correlation with the average annual temperature of their region of origin. Thus, thioglucose-acylated GSLs emerge as promising metabolic markers, and possible effectors, of *A. thaliana* seed responses and adaptation to ET.

Our model regarding the accumulation of thioglucose-acylated GSL pathways upon ET in *A. thaliana* seeds is summarized in **Figure 6**. We have previously shown that short-chain and long-chain methionine derived GSLs accumulated in the SE or in SC, respectively (Barreda et al. 2025b). Interestingly, all the thioglucose-acylated GSLs accumulated exclusively in the SE, including the long-chain 8C thioglucose-acylated GSLs for which their non acylated form is accumulated in the SC (**Figure 3b; Supplemental Figure S7**). This specific localization suggested that thioglucose acylation may enable the accumulation of long-chain GSLs in the embryo, an otherwise restricted compartment for their non-acylated counterparts. The molecular basis of this differential compartmentalization remains to be established. One plausible mechanism involves the UMAMIT-GTR transporter cascade that controls GSL seed loading. The initial discovery of GTR1 and GTR2 as essential GSL transporters (Nour-Eldin et al. 2012) and subsequent characterization of the UMAMIT-GTR cascade operating in the funiculus and seed coat (Xu et al. 2023; Sanden et al. 2024) have established the framework for understanding GSL compartmentalization in seeds. Whether these transporters can discriminate between acylated and non-acylated GSL substrates, and thus whether thioglucose acylation itself determines embryo routing independently of temperature, remains an open question. Alternatively, thioglucose acylation may alter the physicochemical properties of GSLs, such as polarity or membrane permeability, in ways that favour accumulation in the embryo independently of active transport.

**Figure 6:**
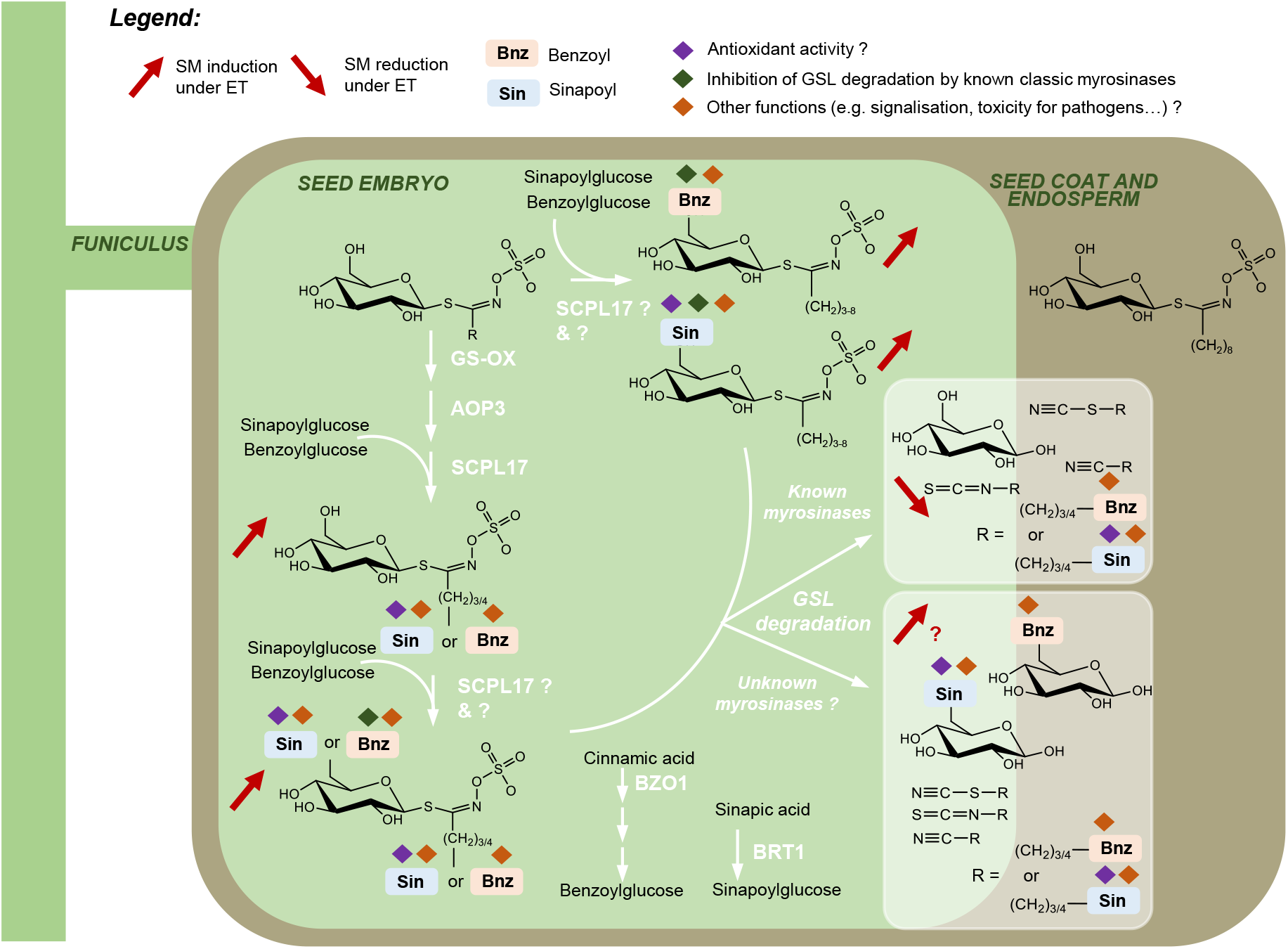
Schematic representation of elevated temperature effect on glucosinolate accumulation in *Arabidopsis thaliana* seeds. Roles are proposed for the sinapoylation and benzoylation occurring in glucosinolate side-chain and thioglucose. Gene acronyms: AOP3, alkenyl hydroxalkyl-producing 3; SCPL17, serine carboxypeptidase-like 17; BZO1, benzoyloxy GSL 1; BRT1, bright trichomes 1.

Different phenotypes were observed in the mutants affected in GSL modification analysed in this study. The reduced seed number per silique observed in aop3-1, bzo1 and scpl17 mutants under ET, but not in wild type, suggests that the acylated GSL pathway contributes to maintaining seed set under ET. This effect could be mediated directly by the acylated GSLs themselves, for instance through a signalling or protective role, or indirectly through the broader transcriptomic and metabolic changes associated with their loss. The normal phenotype of wild type plants under these conditions (27°C/24°C) suggests that the temperature applied is sub-threshold for them, and that the mutant phenotype reflects an increased sensitivity to ET impacting on seed set (**Supplemental Figure S13b**). While no difference in oil accumulation was observed between genotypes, the increased C18:2/C18:3 ratio, known to be temperature-sensitive under ET conditions (Menard et al. 2017) validated the impact of the treatment (**Supplemental Figure S13c**).

Previous studies have established that the acyltransferase SCPL17 (SERINE CARBOXYPEPTIDASE-LIKE 17) mediates the sinapoylation and benzoylation of GSL side-chains, and that BZO1 (BENZOYLOXYGLUCOSINOLATE 1) is essential for the production of side-chain benzoylated GSLs (Kliebenstein et al. 2007; Lee et al. 2012). Building on this knowledge, our work demonstrated that SCPL17 is essential for the sinapoylation and benzoylation of GSL thioglucose moieties, and that corresponding benzoyl groups originate from a benzoyl donor that require BZO1 activity for its synthesis. Hence, SCPL17 and BZO1 are involved in both GSL side-chain and thioglucose acylation. While BAHD acyltransferases catalyse the acylation of several SMs using a CoA esters as activated acyl donors, SCPL enzymes have been described to rather use glucose esters (Ciarkowska et al. 2019; Naake et al. 2024). Therefore, it is most likely that SCPL17 uses both sinapoylglucose and benzoylglucose as activated acyl donors. However, the possibility that SCPL17 may also use sinapoyl CoA and/or benzoyl CoA cannot completely be ruled out without assessing SCPL17 activity.

GSL degradation by myrosinases implies hydrolysis of glucose moieties to form aglycones, which are then converted into bioactive products such as ITCs, thiocyanates, epithionitriles and nitriles involved in biotic stress defences. The question therefore arises whether sinapoyl and benzoyl groups are hydrolysed from GSLs thioglucose moieties prior to GSL degradation or whether myrosinases are able to use thioglucose-acylated GSLs. While some works have suggested that the myrosinases do not accept acylated GSLs as substrates (Rask et al. 2000; Chhajed et al. 2019), likely based on the limited number of known thioglucose-acylated GSLs, these assumptions may primarily apply to side-chain acylated GSLs. Moreover, Reichelt et al. (2002) have suggested that the 6’-benzoyl GSLs from *A. thaliana* Col-0 seeds can be hydrolysed by a commercial myrosinase from *Sinapis alba*.

In conclusion, the newly discovered thioglucose-acylated GSLs produced by SCPL17 and BZO1 accumulate at high levels in *A. thaliana* seeds under ET, contributing both to metabolic responses to ET stress and to adaptation, as reflected by their constitutive accumulation in accessions from diverse geographical origins. Unravelling the specific roles of these GSLs in seeds, alongside assessing their impact on nutritional value for both animals and humans, will be pivotal for safeguarding seed physiological integrity and nutritional quality in a warming climate.

## MATERIALS AND METHODS

### Plant genetic material

When not specified, *A. thaliana* (thale cress) wild-type and transgenic lines were in the Columbia-0 (Col-0) background. The T-DNA insertion mutants of AOP3 (two independent lines, SALK_001655C and SALK_022752C, named *aop3-1* and *aop3-2*, respectively), BZO1 (two independent lines, SALK_094196 and GABI_565B09, named as *bzo1-4* and *bzo1-6*, respectively) and SCPL17 (two lines independent, SALK_075481 and SALK_126264, named as *scpl17-1* and *scpl17-2*, respectively) were kindly provided by Clint Chapple (Purdue University, USA), or obtained from the Eurasian Arabidopsis Stock Centre (uNASC, University of Nottingham, Nottingham, UK). Moreover, 85 Arabidopsis accessions were used for a specific experiment. Accessions, predominantly of European origin, were selected to represent contrasting environments; corresponding details are provided in **Supplemental Dataset 9a**.

### Plant growth conditions and seed harvest

#### Elevated temperature cultures

Seeds from the different genotypes were sowed directly in fertilized soil. Plants were grown under optimal conditions (« control ») in a growth chamber with long day photoperiod (16h light/21°C - 8h dark/19°C). Watering of plants was made with water. At the onset of flowering (occurring between 5 and 6 weeks after seed sowing), elevated temperature (« ET ») was applied to half of the plants of the corresponding experiment by transferring them to another growth chamber (SANYO™, Thermo Fisher Scientific) displaying warmer temperature conditions (16h of light/27°C - 8h of dark/24°C). The other half of the plants, constituting control plants, remained in the initial growth chamber with control temperature conditions. Depending on the experiment, either three (1-5 plants per replica) or six (1-2 plants per replica) biological replicates were used for each sample condition and genotype. Water content of each pot was carefully followed for both control and stressed plants upon the beginning of ET. Plants were watered each day to maintain plant pot water capacities to 85-90%.

Tagging of flower buds allowed us to identify precisely the developmental stages of siliques and seeds (Barreda et al. 2025a). Wild-type and aop3-1 seeds produced either under control or ET conditions were harvested at both torpedo (harvest at 9 and 5 days after flowering [DAF] for control and ET respectively) and mature-green stage (harvest at 14 and 9 DAF for control and ET respectively) as described previously (Barreda et al. 2025a). Upon silique dissection, harvested seeds were directly frozen in Eppendorf tubes placed in liquid nitrogen and stored at −80°C. Plant watering was progressively reduced to maintain plant pot water capacities to 65 %, and then 50 %, once most of the siliques had reached mature stages and dry seeds were harvested. Homogenization of the seed samples was performed using a mortar and pestle with liquid nitrogen. Seed grinded samples were stored at −80 °C prior metabolomic and transcriptomic analysis. Intact dry seeds were kept at both 4°C and −80°C prior to physiological analysis.

Two rosette leaves of medium size (approximate length = 6 cm) were harvested during the fruiting period from aop3-1 and wild-type lines grown under control and ET conditions. Harvested leaves were directly frozen in Eppendorf tubes placed in liquid nitrogen and stored at −80°C. All the leaves harvested for a same sample during the experiment were mixed and homogenised by grinding with pestle in a mortar with liquid nitrogen.

#### Arabidopsis multi-accessions culture

To compare seed glucosinolate accumulation in *A. thaliana* accessions, 85 accessions (**Supplemental Dataset 9a**) were sowed and grown in a greenhouse with a minimum photoperiod of 13 h ensured by supplementary lighting (with Philips-E40 red spectrum – HPS 400W SONT PIA PLUS bulbs; 120 mE.m-2.s-1). Dry seeds were harvested from 2/3 plants for each accession to perform metabolomic analyses. The precise location of each latitude/longitude coordinates were retrieved from the website https://climatecharts.net/ together with their corresponding elevation level (in meters), average annual temperature (in celsius degrees) and annual precipitation sum (in millimeters) between years 1993 and 2022 (**Supplemental Dataset 9a**).

#### Molecular characterization of T-DNA mutants

Plant genomic DNA flanking the T-DNA borders of the mutants was amplified by PCR (**Supplemental Dataset 11**) and sequenced to confirm the mutations. Homozygous lines were then isolated for further characterization.

### Untargeted metabolomic data acquisition and analysis

#### Extraction of polar and semi-polar specialized metabolites and LC-MS/MS injection

Untargeted metabolomic data acquisitions were performed as previously described (Boutet et al., 2022). Briefly, metabolites were extracted from 5-7 mg of grinded seeds, and 50 mg of grinded leaves, using Methanol:Methyl-tert-butyl:Water (1:3:1) buffer, while polar and semipolar metabolites were separated from the oil and protein fractions using a Methanol:Water (1:3) buffer. Polar and semi-polar metabolite fraction was used for metabolomic analyses. Apigenin (500 ng per sample) was used as internal standard. Untargeted metabolomic data were acquired using a UHPLC system (Ultimate 3000 Thermo) coupled to quadrupole time of flight mass spectrometer (Q-Tof Impact II Bruker Daltonics, Bremen, Germany). A Nucleoshell RP 18 plus reversed-phase column (2x 100 mm, 2.7 µm; Macherey-Nagel) was used for chromatographic separation. Samples were injected in both positive and negative electrospray ionisation (ESI) modes (ESI+ and ESI-).

#### Data processing

ESI+ and ESI-data were processed using MZmine 2.52 software (http://mzmine.github.io/) as previously described (Boutet et al. 2022).

#### Ion annotation

Metabolite annotation was performed in four steps: 1) ‘custom database search’ module from MZmine was used to compare the obtained LC-MS/MS data with the IJPB chemistry and metabolomic platform homemade experimental (m/z absolute tolerance of 0.0025 and RT tolerance of 0.3 min) and exact mass (m/z absolute tolerance of 0.0025 Da or 6 ppm) libraries containing respectively 166 standards or experimental common features (RT, m/z) and 1112 ion known m/z; 2) LC-MS/MS data were also searched against the available MS^2^ spectral libraries (Massbank NA, GNPS Public Spectral Library, NIST14 Tandem, NIH Natural Product and MS-Dial); 3) a molecular network analysis was used to assign not-annotated metabolites to a metabolic category (see the following section and Boutet et al. (2022), Barreda et al. (2025b, 2025a) and Olivon et al. (2018) for further information); 4) Sirius software (https://bio.informatik.uni-jena.de/software/sirius/) was used to assign a putative annotation to metabolic features that were not annotated during the previous steps; 5) Glucosinolate annotation was performed by manual expertise, based on previously published data and further interpretation.

#### Data normalisation

Raw data were normalised on the internal standard (Apigenin) and weight of seeds used for the extraction. Data from the *aop3-1, scpl17-1, scpl17-2, bzo1-4, bzo1-6* and wild-type experiment were also additionally normalized by the median of samples, two modes (ESI + and ESI −) separately, due to the high number of samples injected for this LC-MS/MS run. Regarding the normalization of the seed coat/seed embryo injection, they were first normalized according to the seed tissue proportion (Seed coat and endosperm: 10 % and Seed embryo: 90 %), following data were normalized by the total sum of accumulation per sample in each mode (ESI + and ESI −), multiplied by 1000.

#### Molecular network generation of untargeted metabolomic data

As described previously, molecular networks were generated with MetGem software (Olivon et al. 2018)(https://metgem.github.io) using the .mgf and .csv files obtained with MZmine2 analysis (Boutet et al., 2022). The molecular networks were optimized for each of the ESI+ and ESI-datasets, and different cosine similarity score thresholds were tested. Molecular networks were exported to Cytoscape software (Shannon et al. 2003)(https://cytoscape.org/) to format the metabolic categories.

#### Statistical analyses of metabolomic data

Statistical analyses were performed using Metaboanalyst software (both 5.0 and 6.0 versions) (Pang et al. 2021, 2024). Autoscale and log10 transformation were applied to the normalized data to perform the statistical analyses. In particular, an ANOVA has been conducted to identify differentially accumulated metabolites among seed developmental stages (globular, transition, torpedo, mature-green, bent cotyledon and dry seed) and seed developing conditions (control and ET) from wild-type genotype. Volcano plots were conducted to identify the metabolites induced by ET separately for each genotype (*aop3-1, aop3-2* and wild-type) and seed developmental stage (torpedo, mature-green and dry seed). Linear model analyses were performed to the identify the metabolites affected by the genotype (*aop3-1, scpl17-1, scpl17-2, bzo1-4, bzo1-6* and wild-type) and ET in dry seeds. Differences were considered significant when adjusted p-value ≤ 0.05. Principal component analysis (PCA) was performed with Metaboanalyst software and PC1 and PC2 scores were graphically plotted with R.

The log_2_ fold-change of the ET/Control (ET/Ctr) ratio of total SM accumulation averages (n=3) was determined for each metabolic category at each seed developmental stage (globular, transition, torpedo, mature-green, bent cotyledon and dry seed).

Correlation analysis between GSL accumulation in *A. thaliana* accession seeds and the different origin location parameters were calculated with using Pearson correlation. The country, latitude and longitude origin of each *A. thaliana* accessions were retrieved from the catalogue of *Arabidopsis thaliana* genetic Variation (https://1001genomes.org/; The 1001 Genomes Consortium 2016). The precise location of each latitude/longitude coordinates were retrieved from the website https://climatecharts.net/ together with their corresponding elevation level, average annual temperature and annual precipitation sum between years 1993 and 2022.

### Transcriptomic analysis

#### RNA extraction

RNAs were extracted with the PicoPure™ RNA Isolation Kit (ThermoFisher) following the corresponding protocol with slight modifications. Briefly, 2 to 5 mg of seeds were grinded with some polyvinylpolypyrrolidone (PVPP; phenolic compound absorbing agent) in liquid nitrogen. 100 µL of extraction buffer was added to each sample. Following, samples were vortexed, incubated at 42°C for 30 min (500 rpm) using a ThermoMixer™ C (Eppendorf), and centrifuged (2 min, 3000 g). Supernatants were collected, without picking up the pelleted material, and placed in the pre-conditioned RNA purification columns to be centrifuged (1 min, 16 000 g). The cell extracts obtained were mixed with 100 µL of 70 % ethanol by pipetting up and down. The mixtures were placed into the pre-conditioned purification columns which were centrifuged 2 min at 100 g, to allow RNA binding to the columns, immediately followed by a centrifugation to remove flow-through (30 sec, 16 000 g). The columns were washed by adding 100 µL of wash buffer 1 and centrifugation (1 min, 8 000 g). DNase treatment was made by adding 40 µL of DNase solution mix (10μl DNase I Stock solution + 30μl Buffer RDD; RNase-Free DNase Set, Qiagen) to each column, followed by 15 min of incubation (ambient temperature) and washing of the column (addition of 40 µL of wash buffer 1 and centrifugation 30 sec at 8 000 g). Columns were washed twice with 100 µL of wash buffer 2 (2 min at 8 000 g) and a last centrifugation was made after flow-through waste discard to remove all traces of wash buffer (1 min, 16 000 g). 12 µL of elution buffer were added to each column, which were incubated 1 min at ambient temperature before RNA elution by centrifugations (1 min at 1 000 g followed by 1 min at 16 000 g). Obtained RNAs were stored at −80 °C until sequencing.

#### RNA sequencing

Sequencing of messenger RNAs (mRNAs) was performed by BGI Genomics (Hong Kong, https://www.bgi.com). Prior to cDNA library construction (Eukaryotic Strand-specific Transcriptome), quality and concentrations of mRNAs were checked with the bioanalyzer from BGI platform. The sequencing (paired-end reads of 100 bp, 20M clean reads per sample) was performed with DNA Nanoballs Technology (DNBSEQ™). After sequencing, raw reads were filtered by the BGI platform with the filtering software SOAPnuke (version 1.5.2, https://github.com/BGI-flexlab/SOAPnuke) (Cock et al. 2009; Chen et al. 2018) (Parameters used: l 15 -q 0.2 -n 0.05) to reach a Phred+33 fastq quality score.

#### Annotation and statistical analysis of transcriptomic data

High quality raw reads were aligned to the *A. thaliana* genome (TAIR10.1) using STAR (Dobin et al. 2013). Reads were counted and normalized using the R package DESeq2 (Love et al. 2014). In order to evaluate the individual effects of the seed developmental stage (globular, transition, torpedo, bent cotyledon, mature-green and dry seed), and growing condition (Control and ET) on gene expression, a multifactorial analysis was conducted using the multifactor designs method of the DEseq2 R package (Love et al. 2014)(https://bioconductor.org/packages/release/bioc/html/DESeq2.html). This method evaluates the weight of each factor considered in the analysis and its impact on differentially expressed genes (DEGs), according to a false discovery rate (FDR)-adjusted P-value <0.05. Moreover, pairwise comparisons were performed with the BGI data visualisation and analytical platform “Dr. Tom” (https://eu-biosys.bgi.com/).

### Clustering and heatmap analyses

The R packages “pheatmap”, “tidyverse” and “dendextend” were used to perform hierarchical clustering analysis (ward.D) and create heatmaps using omics data.

### AOP3 gene correlation network with SMs and genes coding for enzymes involved in SM modifications

Untargeted metabolomic and transcriptomic data correlations from the 6 seed developmental stages (globular, transition, torpedo, bent cotyledon, mature-green and dry seed) developed under both control and ET conditions previously published (Barreda et al. 2025a) were used to create AOP3 gene correlation network with SMs and genes coding for enzymes involved in SM modifications (according to Barreda et al. (2024)) by using Cytoscape software (Shannon et al. 2003).

### Physiological analyses

#### Seed number per silique

Four siliques per plant (each plant consisting in one of the 6 biological replicates of a sample (i.e. one genotype line and one condition) were collected to count the number per siliques with a binocular magnifier. Statistical analyses were carried out with a Mann-Whitney tests to identify the differences among each genotype between control and ET conditions. Differences were considered significant when adjusted p-value ≤ 0.05.

#### Plant dry weights

After collecting seeds, plants were placed in a drying room for a few days. Plant vegetative parts were harvested and then incubated at 45°C for 48 h. Plants were directly weighted to keep the plants dry. Statistical analyses were carried out with a Mann-Whitney tests to identify the differences among each genotype between control and ET conditions. Differences were considered significant when adjusted p-value ≤ 0.05.

#### Seed specialized metabolite antioxidant capacity analysis

Specialized metabolites were extracted from 12 mg of dry seeds as presented previously (see ‘Extraction of polar and semi-polar specialized metabolites’) for the 6 biological replicates of each genotype developed either in control or ET conditions. Evaporated SM extracts were stored at −80°C.

DPPH (2,2-diphenyl-1-picrylhydrazyl) radical scavenging assay was performed to evaluate the antioxidant capacities of the extracted SM samples. DPPH is a stable free radical that can be used to measure radical scavenging activities of antioxidants by spectrophotometry. Indeed DPPH colour turns from dark purple to light yellow upon its reduction resulting from the transfer of an hydrogen from an antioxidant molecule (Mishra et al. 2012; Liu et al. 2013; Bilel et al. 2020; Salihović et al. 2022). The following described protocol was settled up to evaluate the antioxidant capacities of *A. thaliana* seed polar and semi polar metabolites extracted, in accordance with previously described procedural recommendations (Mishra et al. 2012).

A DPPH stock solution (1.51 mM) was prepared by dissolving 30 mg of DPPH in 50 mL of 96 % ethanol and put in agitation in obscurity for 4-5 hours. Following, the DPPH solution of use (300 μM) was prepared by diluting DPPH stock solution in 96 % ethanol. The solution of use is sonicated for 15 minutes. The extracted metabolites were solubilised in 1.8 mL of 96% ethanol and sonicated in an ice-batch for 10 minutes. Following a centrifugation was performed to precipitate the possible pellet. Five concentrations (C_1_; C_1/2_; C_1/4_; C_1/8_; C_1/16_) were obtained by serial dilution of ½ made for each sample (900 μL of the precedent solution is diluted in 900 μL of 96 % ethanol).

200 μL of DPPH solution (300μM) were added to 200 μL of each concentration sample in a 96 multiwell plate. Three technical replicates were used for each concentration and samples. Control was made with DPPH and ethanol only and Blank with Ethanol only. The absorbance of the samples was then measured with a spectrophotometer (SPARK Control, V2.3 [®2018, TECAN Austria GmbH]) at 517 nm for 990 min to obtain the stabilized absorbance value of all the samples. To obtain the EC50, which is the efficient concentration required to decrease the initial DPPH concentration by 50%, the radical scavenging activity (RSC) was calculated with the following equation:

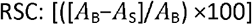

A_B_ and A_S_ correspond to the absorbances at 517 nm of the radical (DPPH) in the absence (Control) and presence of sample extracts respectively at 990 min, where the reaction is in a steady state.

The RSC values were plotted in a graph according to the different concentrations of each biological replicate. Linear regression lines were created for each sample. According to the antioxidant activity of the samples, either C_1_ or C_1/16_ concentrations were not considered to obtain the highest r^2^ coefficient (≥ 0.90). EC50s were calculated with the following equation:

EC50 = 50 / α, where α corresponds to linear coefficient of the linear regression line.

Statistical analyses were carried out with an ANOVA test followed by Newman-Keuls post-hoc test to identify the differences among between genotypes and conditions. Differences were considered significant when adjusted p-value ≤ 0.05.

#### Fatty acid analysis

Total fatty acid analyses were performed as previously described (Li et al. 2006; Kazaz et al. 2020) on pools of *A. thaliana* seeds. Briefly, pools of 50 seeds were incubated in a glass reaction tube in 1.3 ml of methanol/sulfuric acid/toluene (100:2.5:30, v/v/v) at 94°C for 90 min. Fatty acyl methyl esters were then extracted in 0.5 ml of hexane after the addition of 1.5 ml of a 154 mM NaCl solution. After shaking and centrifugation, 1 μl of the upper organic phase was analyzed by gas chromatography (Agilent, 7890B GC system) on a CP-Wax 52 CB 30 m x 530 µm x 1 µm column (Agilent). Fatty acid methyl esters were quantified using 17:0 for calibration. The gas chromatographic acquisition parameters were as follows: hydrogen was used as carrier gas at the rate of 5.25 ml.min^−1^; injector temperature of 250°C; detector temperature of 250°C; oven initial temperature of 175°C for 0 min, followed by a ramp of 0.5°C.min^−1^ to 190°C, then a ramp of 3°C.min^−1^ to 244°C, this final temperature being held for 5 min; splitless injection.

### Statistical analyses

GraphPad Prism® V8.3.0 software, “agricolae” and “car” R packages were used to perform the statistical analysis of the above mentioned physiological and antioxidant capacities analyses.

### Accession numbers

The Arabidopsis Genome Initiative accession numbers for the genes and gene products mentioned in this article are as follows: AT4G03050 (*AOP3*), AT3G12203 (*SCPL17*) and AT1G65880 (*BZO1*).

## SUPPLEMENTAL DATA

**Supplementary Figure 1: Gene ontology enrichment of genes induced by elevated temperature during seed development**. Top-20 gene ontologies are shown for the subsets of genes induced by elevated temperature at the following seed developmental stages: globular, transition, torpedo, bent cotyledon, mature green and dry seed. The enrichment analyses were performed with Shinygo 8.0 software (http://bioinformatics.sdstate.edu/go/)(Ge et al., 2020).

**Supplementary Figure 2: Elevated temperature differentially expressed genes from major metabolic pathways**. Gene expression heatmap are showed for the elevated temperature differentially expressed genes belonging to a) flavonoids, b) glucosinolates and c) cinnamic acids and derivatives pathways. The Arabidopsis Genome Initiative code (AGI code), acronym, functions, average read counts and log_2_(ET/Control of average total expression) are showed for each gene.

**Supplementary Figure 3: Principal component analysis performed on untargeted metabolomic data obtained from torpedo, mature-green and dry seeds of both *aop3-1* mutant and wild-type genotypes developed under control or elevated temperature**.

**Supplementary Figure 4: Volcano plot analyses performed to identify the specialized metabolites affected by elevated temperature in torpedo, mature-green and dry seeds of both *aop3-1* mutant and wild-type genotypes developed under control or elevated temperature**. Specialized metabolites with log2[ET/Ctr] ≤ −1 and log2[ET/Ctr] ≥ 1 were considered as repressed and induced by elevated temperature respectively. The number of induced and repressed metabolites belonging to known metabolic categories are given for each genotype and seed developmental stage.

**Supplementary Figure 5: Glucosinolates affected by elevated temperature in *aop3-1* mutant and wild-type at dry seed developmental stage**. Relative accumulation heatmaps, log_2_([elevated temperature/control] average total accumulation) (log_2_[ET/Ctr]), control accumulation averages, metabolite identity (ID), mass/charge ratio (*m/z*), retention time (RT), putative annotation and spectral annotation are listed for each glucosinolate.

**Supplementary Figure 6: Heatmap displaying the relative accumulations of the differentially accumulated glucosinolate in the different seed developmental stages and leaves in *aop3-1* mutant and wild-type genotypes developed under control or elevated temperature**. The metabolite identity (ID), mass/charge ratio (*m/z*), retention time (RT) and putative annotation are given for each glucosinolate and related degradation products.

**Supplementary Figure 7: Distribution of glucosinolate and related degradation products between Seed coat & endosperm (SC) and seed embryo (SE) in *aop3-1* mutant and wild-type dry seeds developed under control and elevated temperature**. Metabolite identity (ID), mass/charge ratio (*m/z*), retention time (RT), putative annotation are listed for each glucosinolate and degradation product. Abbreviations of glucosinolate (GSL) and isothiocyanate (ITC) names: 3MSOP GSL, 3-methylsulfinylpropyl GSL; 3SIN GSL, 3-sinapoyloxypropyl GSL; 4MSOB GSL, 4-methylsufinylbutyl GSL; 4MSOB ITC, 4-methylsufinylbutyl ITC; 4MTB GSL, 4-methylthiobutyl GSL; 4SIN GSL, 4-sinapoyloxybutyl GSL; 5BZO GSL, 5-benzoyloxypentyl GSL; 6’-sin-3BZO GSL, 6’-sinapoyl-3-benzoyloxypropyl GSL; 6’-sin-4BZO GSL, 6’-sinapoyl-4-benzoyloxybutyl GSL; 6’-bnz-4BZO GSL, 6’benzoyl-4-benzoyloxybutyl GSL; 6’-bnz-4-MSOB GSL, 6’-benzoyl-4-methylsulfinylbutyl GSL; 6’-bnz-5MTP GSL, 6’-benzoyl-5-methylthiopentyl GSL; 6’-bnz-6MTH GSL, 6’-benzoyl-6-methylthiohexyl GSL; 6’-bnz-7MTH GSL, 6’-benzoyl-7-methylthioheptyl GSL; 6’-bnz-8MSOO GSL, 6’-benzoyl-8-methylsulfinyloctyl GSL; 6’bnz-8MTO GSL, 6’-benzoyl-8-methylthiooctyl GSL; 6’-bnz-indol-3-ylmethyl GSL, 6’-benzoyl-indol-3-ylmethyl GSL; 6-MTH GSL, 6-methylthiohexyl GSL; 6MTH ITC, 6-methylsufinylhexyl ITC; 6’-sin-3SIN GSL, 6’-sinapoyl-3-sinapoyloxypropyl GSL; 6’-sin-4-MSOB GSL, 6’-benzoyl-4-methylsulfinyl GSL; 6’-sin-4MSOB GSL, 6’-sinapoyl-4-methylsulfinylbutyl GSL; 6’-sin-4-MTB GSL, 6’-benzoyl-4-methylthiobutyl GSL; 6’-sin-5MTP GSL, 6’-sinapoyl-5-methylthiopentyl GSL; 6’-sin-6MTH GSL, 6’-sinapoyl-6-methylthiohexyl GSL; 6’-sin-7MTH GSL, 6’-sinapoyl-7-methylthioheptyl GSL; 6’-sin-7MTH GSL, 6’-sinapoyl-7-methylsufinylheptyl GSL; 6’-sin-8MSOO GSL, 6’-sinapoyl-8-methylsulfinyloctyl GSL; 6’-sin-8MTO GSL, 6’-sinapoyl-8-methylthiooctyl GSL; 8MSOO GSL, 8-methylsulfinyloctyl GSL; 8MSOO ITC, 8-methylsulfinyloctyl ITC; 9-(MSO)nonanenitrile, 9-methylsufinylnonanenitrile; OH-6MSOH GSL, 6-hydroxy-6-methylsulfinylhexyl GSL; OH-8MSOO GSL, 8-hydroxy-8-methylsulfinyloctyl GSL.

**Supplementary Figure 8: Glucosinolates affected by elevated temperature in *aop3-2* mutant and wild-type at dry seed developmental stage**. Relative accumulation heatmaps, log_2_(elevated temperature/control average total accumulation) (log_2_[ET/Ctr]), control accumulation averages, metabolite identity (ID), mass/charge ratio (*m/z*), retention time (RT), putative annotation and spectral annotation are listed for each glucosinolate.

**Supplementary Figure 9: Volcano plot analyses performed to identify the specialized metabolites affected by elevated temperature in dry seeds of both *aop3-2* mutant and wild-type genotypes developed under control or elevated temperature**. Specialized metabolites with log2[ET/Ctr] ≤ −1 and log2[ET/Ctr] ≥ 1 were considered as repressed and induced by elevated temperature respectively. The number of induced and repressed metabolites belonging to known metabolic categories are given for each genotype and seed developmental stage.

**Supplementary Figure 10: Statistical analyses performed on transcriptomic data obtained from mature-green seed developmental stage of wild-type and *aop3-1* mutant genotypes developed under control or elevated temperature**. a) Principal component analysis. b) Genes differentially expressed (up or down-regulated) upon elevated temperature. c) Venn diagram presenting the genotype-specific and common up-regulated and down-regulated genes.

**Supplementary Figure 11: Effect of elevated temperature on genes coding for proteins involved in glucosinolate degradation and transport**. Heatmaps displaying the expression of differentially expressed genes coding for proteins involved in a) glucosinolate degradation and b) transport.

**Supplementary Figure 12: AOP3, SCPL17, BZO1 and 4CL gene expression atlas**. Data from https://bar.utoronto.ca/eplant (Klepikova et al., 2016).

**Supplementary Figure 13: Phenotypic analysis of *aop3, bzo1* and *scpl17* and wild-type plants and seeds developed under control and elevated temperature conditions.** a) Dry plant weight of *aop3, bzo1* and *scpl17* mutants and wild-type plants developed under control and elevated temperature conditions. Differences among conditions are indicated (Mann-Whitney tests - p-value ≤ 0.05 [*], p-value ≤ 0.01[**]) b) Number of seed per silique for *aop3, bzo1, scpl17* and wild-type genotypes grown under control and elevated temperature conditions. Differences among conditions are indicated (Mann-Whitney tests - p-value ≤ 0.01 [**], p-value ≤ 0.001[***]) c) C18:2/C18:3 ratio in dry seeds of *aop3, bzo1, scpl17* and wild-type lines grown under control and elevated temperature conditions. Differences among conditions are indicated (Kruskal-Wallis test - p-value ≤ 0.05 [*]. d) Antioxidant capacities of polar and semi-polar metabolites extracted from dry seeds of *aop3, bzo1, scpl17* and wild-type genotypes developed under control and elevated temperature conditions. The smallest de EC50 value (effective concentration which scavenges 50% of DPPH [2,2-diphenyl-1-picrylhydrazyl] radical) is, the strongest is the antioxidant capacity. Differences among conditions are indicated (ANOVA test, Newman-Keuls post-hoc test).

**Supplementary Figure 14: Heatmap displaying the relative accumulation of the specialized metabolites that were determined to be affected by elevated temperature and/or genotype**. The number metabolites belonging to known metabolic categories are given for each heatmap cluster (defined according to specialized metabolite accumulation profiles).

**Supplementary Figure 15: Differences of glucosinolate accumulations in aop3, scpl17 and bzo1 mutant compared to wild-type in dry seeds**. The log2(Mutant/WT control accumulations), metabolite identity (ID), mass/charge ratio (*m/z*), retention time (RT) and putative annotation are listed for each glucosinolate.

**Supplemental Dataset 1: Metabolomic analysis (LC-MS/MS) of *Arabidopsis thaliana* Col-0 wild-type seeds at the globular, transition, torpedo, bent cotyledon, mature-green, and dry stages from plants grown under control and elevated temperature conditions. a)** Averages of annotated metabolic category total accumulation. **b)** values of the elevated temperature/control metabolite averages of *Arabidopsis thaliana* globular, transition, torpedo, bent cotyledon, mature-green and dry seeds samples from data previously published in Barreda et al. (2025) Sci. data 12, 306 (DOI:10.1038/s41597-025-04563-2). **c)** Elevated temperature-differentially accumulated metabolites during *Arabidopsis thaliana* seed development. **d)** Top-15 elevated temperature-induced specialized metabolites attributed to a metabolic category.

**Supplemental Dataset 2: Transcriptomic analysis (RNASeq) of *Arabidopsis thaliana* Col-0 wild-type seeds at the globular, transition, torpedo, bent cotyledon, mature-green, and dry stages from plants grown under control and elevated temperature conditions. a)** Elevated temperature-differentially expressed genes during *Arabidopsis thaliana* seed development. **b)** Top-15 elevated temperature-induced genes for each seed developmental stage. **c)** Genes coding for enzymes putatively involved in specialized modifications affected by elevated temperature.

**Supplemental Dataset 3: Metabolomic analysis (LC-MS/MS) of *Arabidopsis thaliana* Col-0 wild-type and *aop3-1* seeds from torpedo, mature-green, and dry stages from plants grown under control and elevated temperature conditions. a)** Annotation and information on *Arabidopsis thaliana aop3-1* and wild-type specialized metabolites from torpedo, mature-green and dry seed stages and leaves developed under control or elevated temperature conditions. **b)** Relative accumulation of *Arabidopsis thaliana aop3-1* and wild-type specialized metabolites from torpedo, mature-green and dry seed stages and leaves developed under control or elevated temperature conditions (LC-MS/MS).

**Supplemental Dataset 4: Glucosinolate accumulation in seed embryo and seed coat & endosperm tissues of *aop3-1* and wild-type genotypes under control or elevated temperature conditions**.

**Supplemental Dataset 5: Metabolomic analysis (LC-MS/MS) of *Arabidopsis thaliana* Col-0 wild-type and *aop3-2* dry seeds from plants grown under control and elevated temperature conditions. a)** Annotation and information on *Arabidopsis thaliana aop3-2* and wild-type dry seed specialized metabolites developed under control or elevated temperature conditions. **b)** Relative accumulation of *Arabidopsis thaliana aop3-2* and wild-type dry seed specialized metabolites developed under control or elevated temperature conditions (LC-MS/MS).

**Supplemental Dataset 6: Transcriptomic analysis of *Arabidopsis thaliana* Col-0 wild-type and *aop3-1* mature-green seeds from plants grown under control and elevated temperature conditions. a)** Transcriptomic data expression of *aop3-1* and wild-type genotypes at mature-green stage developed under control or elevated temperature conditions. **b)** Elevated temperature-differentially expressed genes at mature-green seed developmental stage in *aop3-1* mutant and wild-type. **c)** Elevated temperature - differentially expressed genes at mature-green seed developmental stage in *aop3-1* mutant and wild-type – differences among genotypes. **d)** Log_2_ ([wild-type/*aop3-1*] of gene expression averages) values for control and elevated temperature conditions.

**Supplemental Dataset 7: Fatty acid analysis of *Arabidopsis thaliana* Col-0 wild-type, *aop3, scpl17* and *bzo1* dry seeds from plants grown under control and elevated temperature conditions. a)** Detailed fatty acid composition of seeds from *Arabidopsis thaliana* wild-type (WT), aop3, scpl17 and bzo1 lines produced under control and elevated temperature conditions. **b)** Kruskal-Wallis test results of fatty acid composition of seeds from *Arabidopsis thaliana* wild-type (WT), *aop3, scpl17* and *bzo1* lines developed under control and elevated temperature. **c)** C18:2/C18:3 ratio values from *Arabidopsis thaliana* seeds from wild-type (WT), aop3, scpl17 and bzo1 lines produced under control and elevated temperature conditions.

**Supplemental Dataset 8: Metabolomic analysis (LC-MS/MS) of *Arabidopsis thaliana* Col-0 wild-type, *aop3, scpl17* and *bzo1* dry seeds from plants grown under control and elevated temperature conditions. a)** Annotation and information on *Arabidopsis thaliana aop3-1, bzo1-4, bzo1-6, scpl17-1, scpl17-2* and wild-type specialized metabolites from dry seeds developed under control or elevated temperature conditions. **b)** Relative accumulation of *Arabidopsis thaliana aop3-1, bzo1-4, bzo1-6, scpl17-1, scpl17-2* and wild-type specialized metabolites from dry seeds developed under control or elevated temperature conditions (LC-MS/MS). **c)** Linear model analysis on *Arabidopsis thaliana* dry seed specialized metabolites. d) Specialized metabolites affected by the genotype and/or seed developing condition.

**Supplemental Dataset 9: Analysis of glucosinolate accumulation in *Arabidopsis thaliana* accessions. a)** *Arabidopsis thaliana* accessions used in this study and their localisation origin parameters. **b)** Glucosinolate relative accumulation in Arabidopsis thaliana accession seeds (LC-MS/MS). **c)** Pearson correlations between glucosinolate accumulation in dry seeds from *Arabidopsis thaliana* accessions and corresponding origin parameters: average annual temperature, latitude, longitude, precipitation and elevation. **d)** Kruskal-Wallis results from comparisons between glucosinolate accumulation - origin location parameter average correlations.

**Supplemental Dataset 10: 6’-acyl-thioglucose glucosinolates identified in this study and previous studies**.

**Supplemental Data Set 11: Primer genotyping information**.

## ACKNOWLEDGEMENTS

The authors truly thank Clint Chapple (Purdue University, United States of America) for providing seeds of scpl17 and bzo1 T-DNA insertion mutant lines. They also thank Anne-Solenn Valadon and Chandrodhay Saccaram (Institute Jean-Pierre Bourgin for Plant Sciences – IJPB, INRAE, France) for their help and advice for PCA and heatmap analysis. They are also grateful to Florian Pion and Adithya Raveendran (IJPB, INRAE, France) for their help and advice on DPPH test, Ľubomír Harenčár (Plant Science and Biodiversity Center, Slovak Academy of Sciences, Slovakia) for his help in plant culture, and to Gwendal Cueff (IJPB, INRAE, France; new address: UNH, Université de Clermont Auvergne, France) and Gabrièle Adam (IJPB, INRAE, France; new address: IPS2, Université Paris-Saclay, France) for the development, help and advice about the R script for metabolomic data processing.

The PhD scholarship of L.B. was funded by the ‘Plant biology and breeding’ (BAP) department of INRAE (“Appel à projets scientifiques BAP 2021” to M.C.) and by the project ‘Seed MetSpe’ of the Labex Saclay Plant Sciences-SPS (ANR-10-LABX-0040-SPS to L.L. and M.C.). This work was supported by the ‘Plant biology and breeding’ (BAP) of INRAE (grant “Appel à projets scientifiques BAP 2020” to M.C.), by the project ‘Seed MetSpe’ of the Labex Saclay Plant Sciences-SPS (ANR-10-LABX-0040-SPS to L.L. and M.C.), by the project ‘SeedNapic’ of the Labex Saclay Plant Sciences-SPS (ANR-10-LABX-0040-SPS to M.C.), and by a project financed by Plant2Pro Carnot institute (‘BrassiMet’ to M.C.). This work also benefitted of the financial support of the French government through the National Research Agency (ANR) as part of ‘Growing and Protecting crops Differently’ French Priority Research Program (PPR-CPA) in the framework of SUCSEED project (ANR-20-PCPA-0009, to L.R. and M.C.), and as part of France 2030 in the framework of OLEOPROTID project (ANR-23-DIVP-0004, to L.R., L.L. and M.C.). The IJPB benefits from the support of Saclay Plant Sciences-SPS (ANR-17-EUR-0007). This work has benefited from the support of IJPB’s Plant Observatory platforms PO-Chem and PO-Plants.

## DATA AVAILABILITY

Untargeted metabolomic raw data (.mzXML) for both negative and positive ESI modes, and metadata have been deposited at the MassiVE data repository portal with the following identifiers:

- Specialized metabolome data of developing seeds from wild-type and *aop3* genotypes developed under elevated temperature – MassIVE MSV000099595 (doi:10.25345/C52F7K460)
- Specialized metabolome data of dry seeds from wild-type and *aop3* genotypes developed under elevated temperature – MassIVE MSV000099412 (doi:10.25345/C5FN11542)
- Specialized metabolome data of dry seeds from wild-type, *aop3, scpl17* and *bzo1* genotypes developed under elevated temperature - MassIVE MSV000099408 (doi:10.25345/C5TT4G626)
- Seed embryo and seed coat/endosperm specialized metabolome of *Arabidopsis thaliana* wild-type and *aop3* mutant seeds developed under elevated temperature - MassIVE MSV000099458 (doi:10.25345/C5RR1Q10F)
- Specialized metabolome data from Arabidopsis accession seeds (ESI −) – MassIVE MSV000099593 (doi:10.25345/C59Z90R45)

The transcriptomic RNA-Seq raw data (FASTQ) have been deposited at the National Center for Biotechnology Information (NCBI) Transcriptome Shotgun Assembly Sequence Database (TSA) with BioProject identification PRJNA1344327.

Reviewers link:

https://dataview.ncbi.nlm.nih.gov/object/PRJNA1344327?reviewer=6ihcbu0u7egjcgc10suvtsn11t

## AUTHOR CONTRIBUTIONS

MC directed the research. MC designed the research, with the help of LL and LB. LB, MLN, CBr, DDV, CBo, DG, NK and MC grew the plants and harvested seeds. LB, DDV and MC performed mutants genotyping. MLN and EF dissected and harvested seed coats/endosperms and seed embryos. LB, MLN and NK performed metabolite extractions from seeds and leaves. LB, SBo, NK, IB, ALC and FP generated the LC-MS/MS data and performed the initial quality analyses on the metabolomic data. LB, SBo, FP, NK and MC performed the metabolites annotation with MetGem, SIRIUS and using MS^2^ spectra. LB performed RNA extractions from seeds. LB and MLN performed the physiological analyses (plant weight, seed number per silique and antioxidant capacity analyses). CA and SBa performed fatty acid analysis. LB, IB, ALC, LR, LL and MC analysed and interpreted the untargeted metabolomic, transcriptomic and physiological data. LB generated all figures and supplemental material. LB and MC wrote the paper, which was edited by LL, LR, SBa and FP.

