## Supplemental Figures for "Elevated temperature drives the biosynthesis of novel acylated glucosinolates in *Arabidopsis thaliana* seeds"

### Globular

| Enrichment FDR | nGenes | Pathway Genes | Fold Enrichment | Pathway |
|---|---|---|---|---|
| 1,19E-06 | 5 | 17 | 74,1026738 | GO:0051259 Protein complex oligomerization |
| 2,72E-05 | 5 | 40 | 31,49363636 | GO:0006721 Terpenoid metabolic process |
| 1,11E-05 | 6 | 59 | 25,62194145 | GO:0010286 Heat acclimation |
| 0,000210286 | 5 | 64 | 19,68352273 | GO:0042542 Response to hydrogen peroxide |
| 2,17E-05 | 7 | 114 | 15,47055821 | GO:0051082 Unfolded protein binding |
| 0,000210286 | 6 | 111 | 13,61886978 | GO:0007623 Circadian rhythm |
| 7,50E-05 | 7 | 143 | 12,33317228 | GO:0006457 Protein folding |
| 1,11E-05 | 9 | 199 | 11,3946825 | GO:0009408 Response to heat |
| 0,00053948 | 8 | 276 | 7,3028722 | GO:0019752 Carboxylic acid metabolic process |
| 2,02E-06 | 14 | 486 | 7,257792742 | GO:0048316 Seed development |
| 2,17E-05 | 13 | 544 | 6,020842246 | GO:0003677 DNA binding |
| 2,17E-05 | 13 | 549 | 5,966007617 | GO:0009651 Response to salt stress |
| 2,17E-05 | 15 | 740 | 5,107076167 | GO:0000976 Transcription cis-regulatory region binding |
| 4,74E-05 | 15 | 810 | 4,665723906 | GO:0009611 Response to wounding |
| 8,41E-07 | 26 | 1629 | 4,021286902 | GO:0003700 DNA-binding transcription factor activity |
| 3,69E-05 | 18 | 1129 | 4,016903132 | GO:0009416 Response to light stimulus |
| 4,98E-06 | 23 | 1485 | 3,902241812 | GO:0006355 Reg. of transcription-DNA-templated |
| 8,58E-06 | 30 | 2485 | 3,041638924 | GO:0005829 Cytosol |
| 1,60E-08 | 49 | 4294 | 2,875059491 | GO:0005515 Protein binding |
| 0,00053948 | 33 | 3744 | 2,220705128 | GO:0005634 Nucleus |

### Bent cotyledon

| Enrichment FDR | nGenes | Pathway Genes | Fold Enrichment | Pathway |
|---|---|---|---|---|
| 3,79E-07 | 5 | 17 | 85,26458282 | GO:0051259 Protein complex oligomerization |
| 0,001307923 | 3 | 17 | 51,15874969 | GO:0034620 Cellular response to unfolded protein |
| 0,001832932 | 3 | 21 | 41,41422594 | GO:0051787 Misfolded protein binding |
| 0,00186322 | 3 | 22 | 39,53176113 | GO:0031072 Heat shock protein binding |
| 0,00027661 | 4 | 32 | 36,2374477 | GO:0042026 Protein refolding |
| 6,62E-06 | 6 | 59 | 29,48131338 | GO:0010286 Heat acclimation |
| 0,001289963 | 4 | 48 | 24,15829847 | GO:0051085 Chaperone cofactor-dependent protein refolding |
| 5,07E-08 | 9 | 114 | 22,88680907 | GO:0051082 Unfolded protein binding |
| 0,000204541 | 5 | 64 | 22,64840481 | GO:0042542 Response to hydrogen peroxide |
| 0,001584729 | 4 | 55 | 21,08360593 | GO:0080044 Quercetin 7-O-glucosyltransferase activity |
| 2,55E-08 | 10 | 143 | 20,27269801 | GO:0006457 Protein folding |
| 1,79E-10 | 13 | 199 | 18,93816362 | GO:0009408 Response to heat |
| 0,002016968 | 4 | 64 | 18,11872385 | GO:0080043 Quercetin 3-O-glucosyltransferase activity |
| 0,00186322 | 5 | 119 | 12,18065469 | GO:0043621 Protein self-association |
| 0,000178274 | 9 | 325 | 8,027988413 | GO:0005506 Iron ion binding |
| 0,001443043 | 7 | 257 | 7,896097553 | GO:0016887 ATP hydrolysis activity |
| 0,000112112 | 11 | 486 | 6,561513164 | GO:0048316 Seed development |
| 0,001832932 | 8 | 375 | 6,184524407 | GO:0020037 Heme binding |
| 0,001443043 | 9 | 454 | 5,746908005 | GO:0006952 Defense response |
| 0,00186322 | 13 | 1022 | 3,687568063 | GO:0007165 Signal transduction |

### Transition

| Enrichment FDR | nGenes | Pathway Genes | Fold Enrichment | Pathway |
|---|---|---|---|---|
| 1,57E-05 | 5 | 17 | 37,0513369 | GO:0051259 Protein complex oligomerization |
| 3,14E-05 | 6 | 38 | 19,8907177 | GO:0048544 Recognition of pollen |
| 3,94E-09 | 14 | 143 | 12,33317228 | GO:0006457 Protein folding |
| 2,92E-07 | 11 | 114 | 12,1554386 | GO:0051082 Unfolded protein binding |
| 3,14E-05 | 8 | 86 | 11,71856237 | GO:0045087 Innate immune response |
| 1,96E-08 | 15 | 199 | 9,495568753 | GO:0009408 Response to heat |
| 3,75E-07 | 15 | 257 | 7,352599929 | GO:0016887 ATP hydrolysis activity |
| 3,94E-09 | 23 | 486 | 5,961758324 | GO:0048316 Seed development |
| 1,74E-05 | 17 | 459 | 4,665723906 | GO:0016757 Glycosyltransferase activity |
| 3,14E-05 | 18 | 544 | 4,168275401 | GO:0003677 DNA binding |
| 3,14E-05 | 19 | 602 | 3,975940803 | GO:0003723 RNA binding |
| 4,68E-06 | 23 | 740 | 3,915425061 | GO:0000976 Transcription cis-regulatory region binding |
| 3,14E-05 | 24 | 922 | 3,279163873 | GO:0009737 Response to abscisic acid |
| 3,29E-05 | 24 | 927 | 3,261476905 | GO:0006468 Protein phosphorylation |
| 1,70E-05 | 27 | 1071 | 3,175828877 | GO:0009507 Chloroplast |
| 2,58E-07 | 39 | 1629 | 3,015965177 | GO:0003700 DNA-binding transcription factor activity |
| 5,37E-06 | 34 | 1485 | 2,884265687 | GO:0006355 Reg. of transcription-DNA-templated |
| 1,83E-07 | 43 | 1885 | 2,873689896 | GO:0005886 Plasma membrane |
| 1,19E-14 | 84 | 3744 | 2,826351981 | GO:0005634 Nucleus |
| 2,29E-14 | 90 | 4294 | 2,640360757 | GO:0005515 Protein binding |

### Mature-green

| Enrichment FDR | nGenes | Pathway Genes | Fold Enrichment | Pathway |
|---|---|---|---|---|
| 2,11E-05 | 5 | 17 | 38,23308686 | GO:0051259 Protein complex oligomerization |
| 4,39E-05 | 6 | 38 | 20,52513084 | GO:0048544 Recognition of pollen |
| 3,60E-06 | 10 | 114 | 11,40285047 | GO:0051082 Unfolded protein binding |
| 7,76E-05 | 8 | 102 | 10,19548983 | GO:0006364 RRNA processing |
| 3,40E-06 | 11 | 143 | 9,999422716 | GO:0006457 Protein folding |
| 4,36E-05 | 9 | 119 | 9,831365191 | GO:0043621 Protein self-association |
| 5,35E-05 | 10 | 162 | 8,024228106 | GO:0016705 Oxidoreductase activity-acting on paired donors-with with incorporation or reduction of molecular oxygen |
| 0,000189471 | 10 | 199 | 6,532286196 | GO:0009408 Response to heat |
| 6,43E-05 | 12 | 257 | 6,069688497 | GO:0016887 ATP hydrolysis activity |
| 1,03E-07 | 21 | 486 | 5,616959674 | GO:0048316 Seed development |
| 3,75E-05 | 14 | 325 | 5,599676721 | GO:0005506 Iron ion binding |
| 0,000187914 | 15 | 459 | 4,248120762 | GO:0016757 Glycosyltransferase activity |
| 5,96E-08 | 32 | 1071 | 3,883996125 | GO:0009507 Chloroplast |
| 6,18E-05 | 19 | 641 | 3,853131686 | GO:0004674 Protein serine/threonine kinase activity |
| 7,73E-05 | 23 | 927 | 3,225272268 | GO:0006468 Protein phosphorylation |
| 6,75E-05 | 26 | 1129 | 2,993626996 | GO:0009416 Response to light stimulus |
| 5,35E-05 | 33 | 1629 | 2,633365467 | GO:0003700 DNA-binding transcription factor activity |
| 0,000121674 | 30 | 1485 | 2,626111016 | GO:0006355 Reg. of transcription-DNA-templated |
| 7,28E-10 | 79 | 4294 | 2,391571292 | GO:0005515 Protein binding |
| 1,03E-07 | 66 | 3744 | 2,291534372 | GO:0005634 Nucleus |

### Torpedo

| Enrichment FDR | nGenes | Pathway Genes | Fold Enrichment | Pathway |
|---|---|---|---|---|
| 0,005706794 | 2 | 6 | 93,50337382 | GO:0001216 DNA-binding transcription activator activity |
| 3,67E-07 | 5 | 17 | 82,5029769 | GO:0051259 Protein complex oligomerization |
| 3,67E-07 | 7 | 59 | 33,28086187 | GO:0010286 Heat acclimation |
| 0,000602288 | 4 | 40 | 28,05101215 | GO:0006721 Terpenoid metabolic process |
| 0,005706794 | 3 | 32 | 26,29782389 | GO:0016102 Diterpenoid biosynthetic process |
| 0,005706794 | 3 | 32 | 26,29782389 | GO:0042026 Protein refolding |
| 0,005706794 | 3 | 32 | 26,29782389 | GO:0010333 Terpene synthase activity |
| 0,000179145 | 5 | 64 | 21,91485324 | GO:0042542 Response to hydrogen peroxide |
| 8,56E-07 | 8 | 114 | 19,6849208 | GO:0051082 Unfolded protein binding |
| 3,67E-07 | 9 | 143 | 17,65448317 | GO:0006457 Protein folding |
| 5,70E-09 | 12 | 199 | 16,9151832 | GO:0009408 Response to heat |
| 0,002875117 | 5 | 119 | 11,78613956 | GO:0043621 Protein self-association |
| 0,004377224 | 5 | 132 | 10,62538339 | GO:0000287 Magnesium ion binding |
| 3,67E-07 | 14 | 486 | 8,080538478 | GO:0048316 Seed development |
| 0,000616931 | 8 | 301 | 7,45541851 | GO:0045893 Pos. reg. of transcription-DNA-templated |
| 5,76E-05 | 12 | 549 | 6,131368775 | GO:0009651 Response to salt stress |
| 0,005706794 | 7 | 321 | 6,117043147 | GO:0055085 Transmembrane transport |
| 0,00011244 | 20 | 1629 | 3,443954837 | GO:0003700 DNA-binding transcription factor activity |
| 2,69E-05 | 27 | 2485 | 3,047796088 | GO:0005829 Cytosol |
| 0,000106359 | 36 | 4294 | 2,351738326 | GO:0005515 Protein binding |

**Supplementary figure 1: Gene ontology enrichment of genes induced by elevated temperature during seed development**. Top- 20 gene ontologies are shown for the subsets of genes induced by elevated temperature at the following seed developmental stages: globular, transition, torpedo, bent cotyledon, mature-green and dry seed. The enrichment analyses were performed with Shinygo 8.0 software (http://bioinformatics.sdstate.edu/go/)(Ge et al., 2020).

### a - Differentially expressed genes (DEGs) from flavonoids pathway

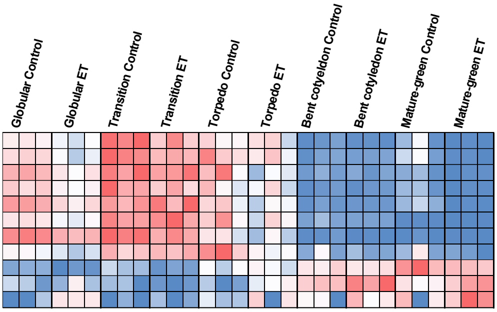


| AGI code | Acronym | Function | Average read counts | Globular | Transition | Torpedo | Bent cotyledon | Mature-green |
|---|---|---|---|---|---|---|---|---|
| AT5G13930 | CHS | Core structure synthesis | *80806.627* | -0.50 | -0.62 | 0.01 | -0.45 | -2.33 |
| AT5G07990 | F3'H (CYP75B1) | Core structure synthesis | *19394.807* | -0.58 | -0.35 | -0.36 | 0.08 | -1.52 |
| AT5G05270 | CHI-L1 | Core structure synthesis | *10777.926* | -0.45 | -0.07 | -0.91 | -0.07 | -1.26 |
| AT1G65060 | 4CL3 | Core structure synthesis | *7363.432* | -0.62 | -0.51 | -0.29 | 0.03 | -2.15 |
| AT3G55120 | CHI | Core structure synthesis | *4680.723* | -0.55 | 0.17 | -0.53 | -0.04 | -2.09 |
| AT5G17220 | GSTF12 | Transport | *3574.619* | -0.34 | 0.08 | -0.49 | -0.49 | -1.30 |
| AT3G59030 | TT12 | Transport | *1781.091* | -0.56 | -0.46 | -1.03 | -2.11 | -1.94 |
| AT4G25640 | DTX35 | Transport | *2446.925* | -0.33 | 0.12 | -1.01 | -0.29 | -0.73 |
| AT5G54160 | OMT1 | Modification | *4181.342* | -0.80 | -0.96 | 0.27 | -0.06 | -0.69 |
| AT5G49330 | MYB111 | TF | *55.168* | 1.42 | -1.95 | 2.67 | 0.98 | 2.11 |
| AT1G56650 | PAP1 | TF | *6.682* | 3.96 | 1.13 | 2.02 | 1.87 | 1.66 |

*Minimal gene expression (normalized read counts)* — *Maximal gene expression (normalized read counts)*

-2 ≤ — ≥ 2
*Log$_2$ (ET/Control of average total expression)*

### b - Differentially expressed genes (DEGs) from glucosinolate (GSL) pathway

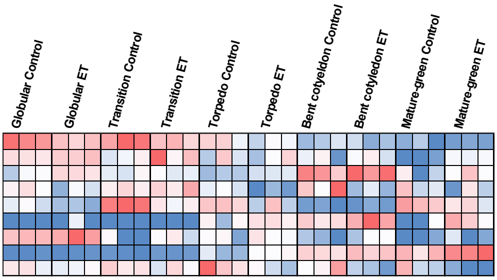


| AGI code | Acronym | Function | Average read counts | Globular | Transition | Torpedo | Bent cotyledon | Mature-green |
|---|---|---|---|---|---|---|---|---|
| AT2G30870 | GSTF10 | Core structure synthesis | *1283.160* | -0.60 | -0.59 | -0.59 | -0.16 | -0.27 |
| AT5G65940 | CHY1 | Cosubstrate pathways | *749.343* | 0.25 | 0.81 | -0.01 | 0.31 | 1.50 |
| AT3G49680 | BCAT3 | Side-Chain Elongation | *848.566* | 0.23 | 0.01 | 0.22 | 0.12 | 0.38 |
| AT3G60140 | BGLU30 | GSL degradation | *135.580* | -0.81 | 0.38 | -2.00 | -1.88 | -0.69 |
| AT3G16390 | NSP3 | GSL degradation | *53.284* | -1.03 | -1.68 | -0.86 | 0.40 | -0.88 |
| AT3G09260 | PYK10 | GSL degradation | *15.613* | | | 1.74 | 1.56 | 5.80 |
| AT4G37400 | CYP81F3 | Side-Chain modification | *13.060* | 0.63 | 1.68 | 0.75 | 1.56 | 2.02 |
| AT4G03050 | AOP3 | Side-Chain modification | *2651.480* | 0.37 | 1.17 | 1.86 | 0.57 | 1.13 |
| AT1G18880 | GTR3 | GSL transport | *107.951* | -0.24 | -0.31 | -1.16 | -1.37 | -2.05 |

*Minimal gene expression (normalized read counts)* — *Maximal gene expression (normalized read counts)*

-2 ≤ — ≥ 2
*Log$_2$ (ET/Control of average total expression)*

### c - Differentially expressed genes (DEGs) from cinnamic acid and derivatives pathway

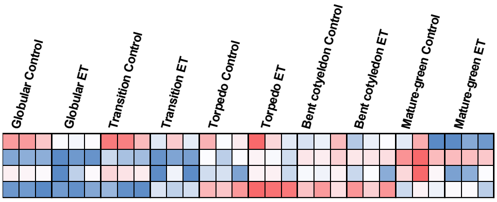


| AGI code | Acronym | Function | Average read counts | Globular | Transition | Torpedo | Bent cotyledon | Mature-green |
|---|---|---|---|---|---|---|---|---|
| AT4G39230 | AT4G39230 | Lignans synthesis and modifications | *862.302* | -0.52 | -0.60 | 0.15 | -0.24 | -1.00 |
| AT5G54160 | OMT1 | Modification | *4181.342* | -0.80 | -0.96 | 0.27 | -0.06 | -0.69 |
| AT5G04230 | PAL3 | Core structure synthesis | *68.326* | -0.86 | -1.95 | 0.82 | -0.50 | -2.58 |
| AT2G23260 | UGT84B1 | Core structure synthesis | *290.717* | 0.27 | 1.28 | 0.66 | 0.30 | -0.08 |

*Minimal gene expression (normalized read counts)* — *Maximal gene expression (normalized read counts)*

-2 ≤ — ≥ 2
*Log$_2$ (ET/Control of average total expression)*

**Supplementary figure 2: Elevated temperature differentially expressed genes from major metabolic pathways.** Gene expression heatmap are showed for the elevated temperature differentially expressed genes belonging to a) flavonoids, b) glucosinolates and c) cinnamic acids and derivatives pathways. The Arabidopsis Genome Initiative code (AGI code), acronym, functions, average read counts and log$_2$(ET/Control of average total expression) are showed for each gene.

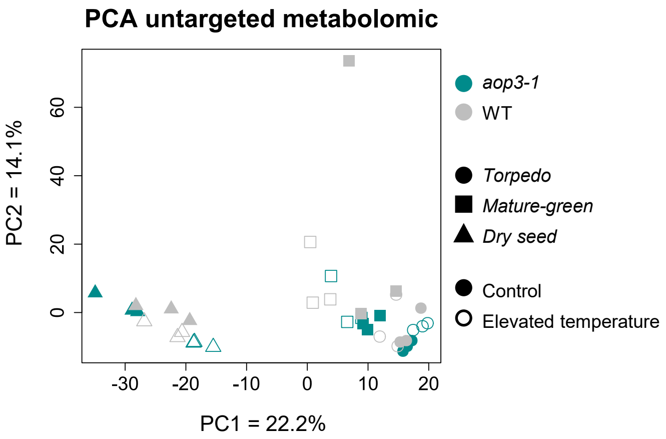


**Supplementary figure 3: Principal component analysis performed on untargeted metabolomic data obtained from torpedo, mature-green and dry seeds of both *aop3-1* mutant and wild-type genotypes developed under control or elevated temperature.**

### Volcano plot torpedo WT

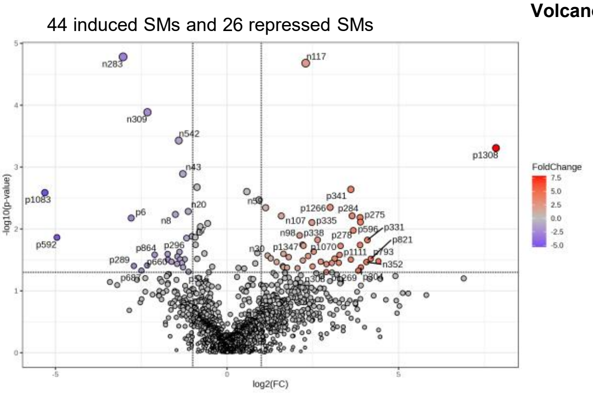


| | WT | |
|---|---|---|
| **Metabolic category** | **Induced** | **Repressed** |
| Cinnamic acid and derivatives | 4 | 2 |
| Lipids and FA | 0 | 2 |
| Carbohydrates | 9 | 1 |
| Glucosinolates | 3 | 1 |
| Amino acids and derivatives | 0 | 1 |
| Benzaldehyde | 0 | 1 |
| Nucleosides and derivatives | 0 | 1 |
| Flavonoids | 4 | 0 |

### Volcano plot torpedo *aop3-1*

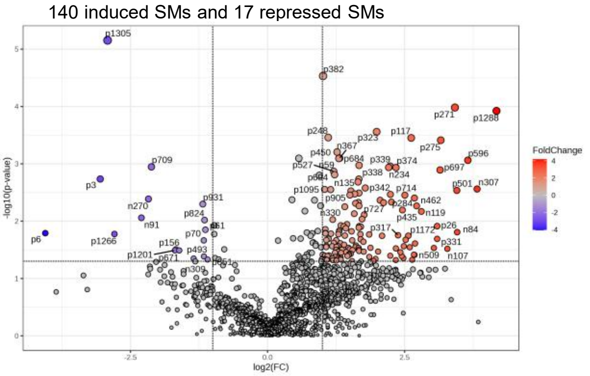


| | *aop3-1* | |
|---|---|---|
| **Metabolic category** | **Induced** | **Repressed** |
| Flavonoids | 11 | 3 |
| Carboxylic acids | 0 | 1 |
| Glucosinolates | 17 | 0 |
| Carbohydrates | 15 | 0 |
| Cinnamic acid and derivatives | 4 | 0 |
| Amino acids and derivatives | 1 | 0 |
| Glucosinolate degradation products | 1 | 0 |
| Lignans | 1 | 0 |
| Terpenoids | 1 | 0 |

### Volcano plot mature-green WT

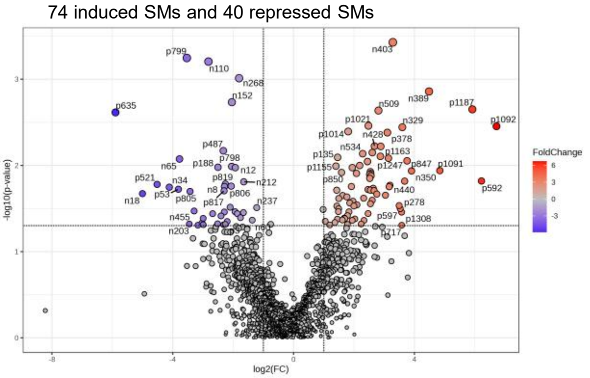


| | WT | |
|---|---|---|
| **Metabolic category** | **Induced** | **Repressed** |
| Cinnamic acid and derivatives | 5 | 4 |
| Flavonoids | 2 | 4 |
| Carbohydrates | 3 | 2 |
| Amino acids and derivatives | 1 | 2 |
| Glucosinolates | 17 | 1 |
| Benzoic acid and derivatives | 0 | 1 |
| Lipids and FA | 2 | 0 |
| Lignans | 1 | 0 |
| Carboxylic acids | 1 | 0 |
| Glucosinolate degradation products | 1 | 0 |
| Benzaldehyde | 1 | 0 |

**Supplementary figure 4: Volcano plot analyses performed to identify the specialized metabolites affected by elevated temperature in torpedo, mature-green and dry seeds of both *aop3-1* mutant and wild-type genotypes developed under control or elevated temperature.** Specialized metabolites with log2[ET/Ctr] ≤ -1 and log2[ET/Ctr] ≥ 1 were considered as repressed and induced by elevated temperature respectively. The number of induced and repressed metabolites belonging to known metabolic categories are given for each genotype and seed developmental stage.

*(Continued 1/2)*

**Volcano plot mature-green *aop3-1***

131 induced SMs and 30 repressed SMs

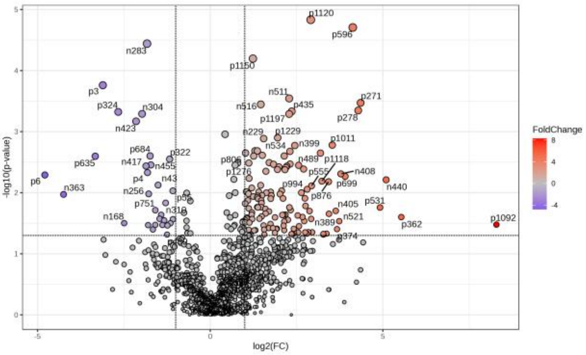


| | *aop3-1* | |
|---|---|---|
| **Metabolic category** | **Induced** | **Repressed** |
| Glucosinolates | 20 | 5 |
| Flavonoids | 6 | 5 |
| Carbohydrates | 8 | 3 |
| Cinnamic acid and derivatives | 5 | 1 |
| Lignans | 0 | 1 |
| Lipids and FA | 2 | 0 |
| Amino acids and derivatives | 1 | 0 |
| Benzaldehyde | 1 | 0 |
| Pyrans | 1 | 0 |

**Volcano plot dry seed WT**

105 induced SMs and 114 repressed SMs

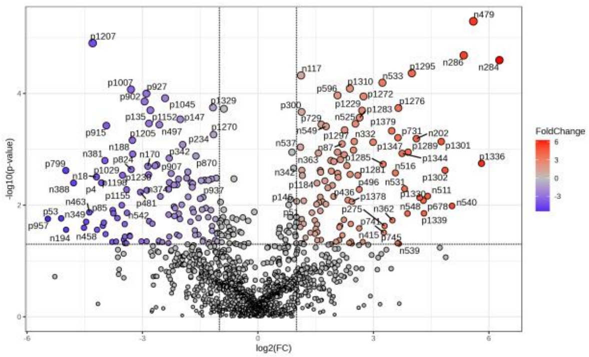


| | WT | |
|---|---|---|
| **Metabolic category** | **Induced** | **Repressed** |
| Flavonoids | 7 | 11 |
| Cinnamic acid and derivatives | 9 | 10 |
| Glucosinolates | 8 | 5 |
| Carbohydrates | 10 | 3 |
| Lipids and FA | 1 | 3 |
| Lignans | 1 | 3 |
| Benzoic acid and derivatives | 0 | 2 |
| Amino acids and derivatives | 1 | 1 |
| Benzaldehyde | 0 | 1 |
| Carboxylic acids | 0 | 1 |
| Benzofuran | 0 | 1 |
| Glucosinolate degradation products | 1 | 0 |
| Choline derivatives | 1 | 0 |
| Phenylethanolamines | 1 | 0 |

**Volcano plot dry seed *aop3-1***

77 induced SMs and 201 repressed SMs

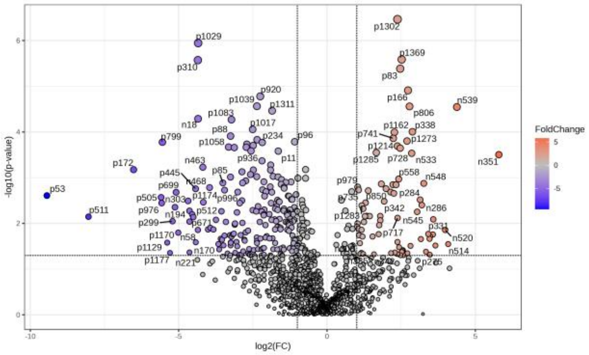


| | *aop3-1* | |
|---|---|---|
| **Metabolic category** | **Induced** | **Repressed** |
| Cinnamic acid and derivatives | 6 | 14 |
| Lipids and FA | 0 | 14 |
| Flavonoids | 6 | 11 |
| Glucosinolates | 8 | 4 |
| Benzaldehyde | 1 | 4 |
| Amino acids and derivatives | 0 | 4 |
| Glucosinolate degradation products | 0 | 4 |
| Lignans | 0 | 3 |
| Carbohydrates | 9 | 2 |
| Carboxylic acids | 0 | 2 |
| Benzoic acid and derivatives | 0 | 1 |
| Terpenoids | 0 | 1 |
| Benzofuran | 0 | 1 |
| Pyrroles | 0 | 1 |
| Nucleosides and derivatives | 1 | 0 |

**Supplementary figure 4: Volcano plot analyses performed to identify the specialized metabolites affected by elevated temperature in torpedo, mature-green and dry seeds of both *aop3-1* mutant and wild-type genotypes developed under control or elevated temperature.** Specialized metabolites with log2[ET/Ctr] ≤ -1 and log2[ET/Ctr] ≥ 1 were considered as repressed and induced by elevated temperature respectively. The number of induced and repressed metabolites belonging to known metabolic categories are given for each genotype and seed developmental stage.

*(Continued 2/2)*

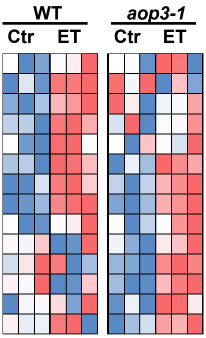


| Log₂(ET/Ctr) WT | Log₂(ET/Ctr) *aop3-1* | Average signal accumulation WT | Average signal accumulation *aop3-1* | ID | *m/z* | RT | Putative annotation | Spectral information |
|---|---|---|---|---|---|---|---|---|
| | 1.57 | *0.628* | *1.075* | n514 | 552.0889 | 3.41 | GSL_n514 | |
| | 2.629 | *0.238* | *0.395* | n520 | 568.0843 | 0.88 | GSL_n520 | |
| | 1.629 | *1.021* | *0.868* | n525 | 640.1203 | 8.55 | 6'-sin-5MTP GSL form 2 | m/z 265; 465; 497 |
| | 4.41 | *2.705* | *0.066* | n537 | 474.1116 | 4.71 | GSL_n537 | |
| | 2.185 | *0.831* | *21.131* | n87 | 422.0257 | 0.75 | 3MSOP GSL | m/z 259; 274 - losses 15,02; 63,99 |
| | 2.63 | *0.144* | *0.466* | p496 | 485.1191 | 3.03 | 4MSB ITC glutathion | |
| 1.245 | 1.621 | *0.812* | *4.260* | n433 | 552.1043 | 11.27 | 6'-bnz-6MTH GSL | m/z 363; 379; 395 |
| 1.832 | 2.532 | *2.719* | *1.656* | n505 | 668.1519 | 11.55 | 6'-sin-7MTH GSL form 2 | m/z 265; 497 |
| 1.11 | 1.104 | *0.538* | *4.405* | n511 | 445.1351 | 3.34 | GSL_n511 | |
| 1.123 | | *23.426* | *19.282* | n335 | 626.1049 | 7.14 | 6'-sin-4MTB GSL form 2 | m/z 265; 465; 481; 497 |
| 5.792 | | *24.512* | *0.128* | n351 | 700.1384 | 10.69 | 6'-sin-4BZO GSL | m/z 265; 465; 481; 497 |
| 2.144 | | *16.132* | *17.355* | n361 | 668.1518 | 10.62 | 6'-sin-7MTH GSL form 1 | m/z 265; 465; 481; 497 |
| 4.084 | | *19.850* | *0.034* | n363 | 510.0750 | 3.99 | GSL_n363 | |
| 3.982 | | *2.956* | *8.175* | n415 | 527.0736 | 7.88 | GSL_n415 | |

*Minimal $log_2(FC)$* — *Maximal $log_2(FC)$*

*Minimal accumulation* — *Maximal accumulation*

**Supplementary figure 5: Glucosinolates affected by elevated temperature in *aop3-1* mutant and wild type at dry seed developmental stage.** Relative accumulation heatmaps, $log_2$([elevated temperature/control] average total accumulation) ($log_2$[ET/Ctr]), control accumulation averages, metabolite identity (ID), mass/charge ratio (*m/z*), retention time (RT), putative annotation and spectral annotation are listed for each glucosinolate.

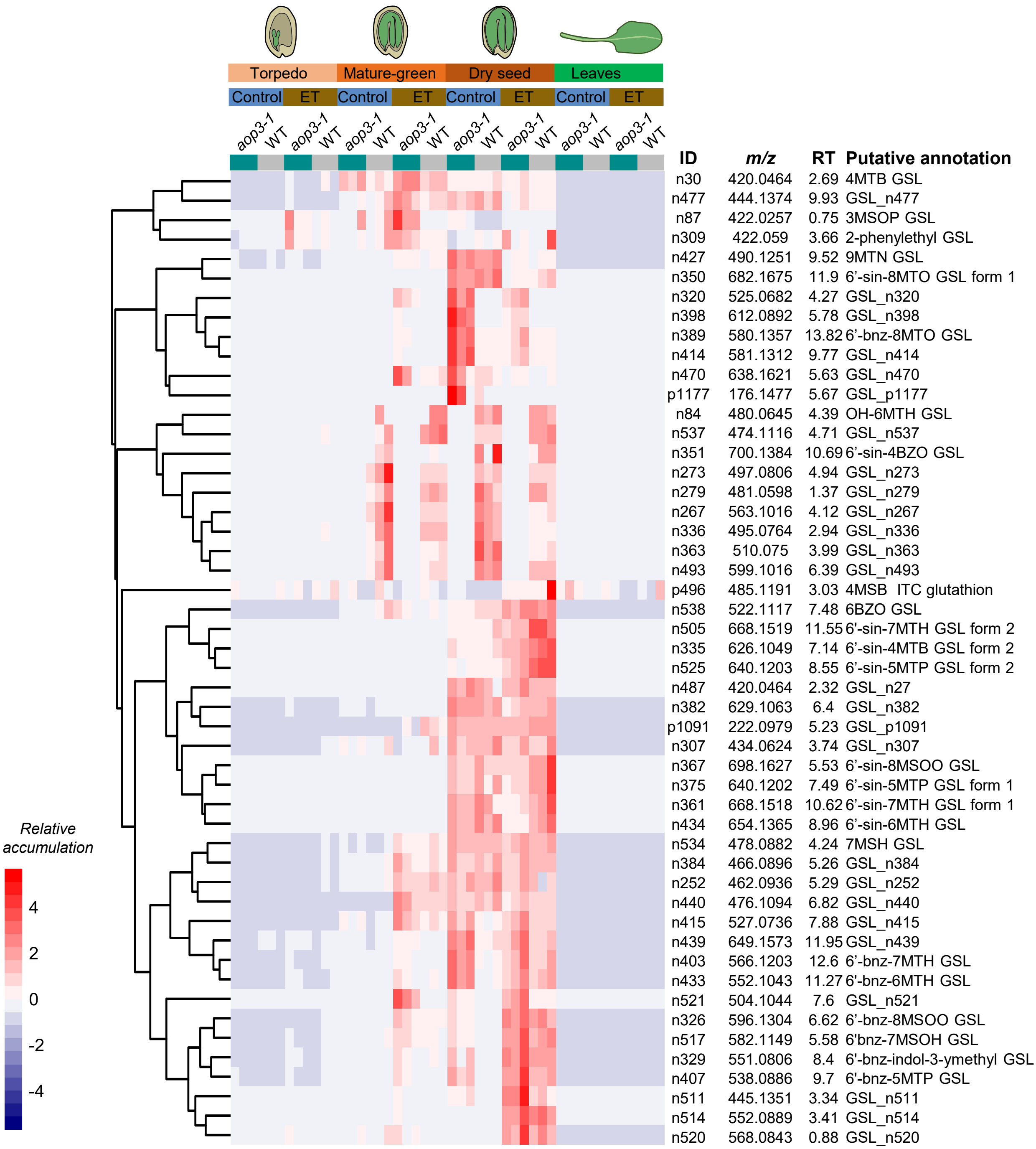


**Supplementary figure 6: Heatmap displaying the relative accumulations of the differentially accumulated glucosinolate in the different seed developmental stages and leaves in *aop3-1* mutant and wild-type genotypes developed under control or elevated temperature.** The metabolite identity (ID), mass/charge ratio (*m/z*), retention time (RT) and putative annotation are given for each glucosinolate and related degradation products

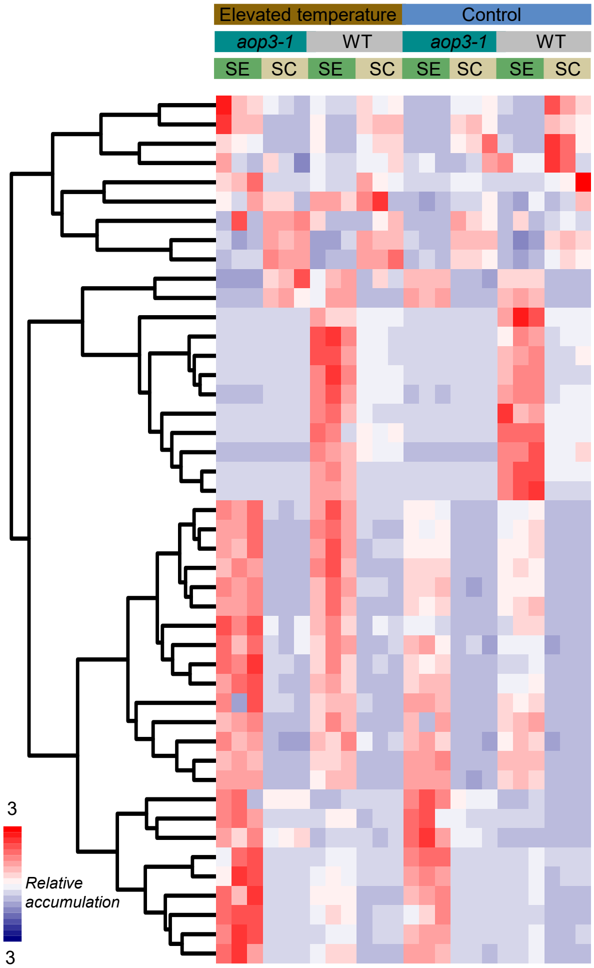


| ID | *m/z* | RT | Annotation |
|---|---|---|---|
| p252 | 206.0667 | 8.21 | 6MTH ITC |
| p169 | 178.0355 | 4.62 | 4MSOB ITC |
| p744 | 234.0981 | 22.54 | 8MSOO ITC form 2 |
| p43 | 234.0981 | 13.25 | 8MSOO ITC form 1 |
| p477 | 485.119 | 3.01 | 4MSOB ITC - glutathion |
| p726 | 449.0676 | 3.2 | Glucobrassicin |
| n5 | 436.0416 | 0.78 | 4MSOB GSL |
| n1 | 492.1041 | 3.51 | 8MSOO GSL |
| n244 | 508.0992 | 3.75 | OH-8-MSOO GSL |
| p31 | 202.1259 | 3.48 | 9-(MSO)nonanenitrile form 1 |
| p3 | 202.1259 | 6.07 | 9-(MSO)nonanenitrile form 2 |
| n236 | 700.1385 | 10.76 | 6'-sin-4BZO GSL form 2 |
| n223 | 686.1228 | 8.58 | 6'-sin-3BZO GSL form 1 |
| n217 | 582.097 | 4.45 | 3SIN GSL |
| n238 | 584.0913 | 10.75 | 6'-bnz-3BZO GSL |
| n241 | 686.1228 | 9.71 | 6'-sin-3BZO GSL form 1 |
| n221 | 700.1383 | 9.8 | 6'-sin-4BZO GSL form 1 |
| n245 | 480.0645 | 4.39 | OH-6MSOH GSL |
| n219 | 596.1122 | 4.9 | 4SIN GSL |
| n231 | 598.1068 | 11.85 | 6'-bnz-4BZO GSL |
| n233 | 508.096 | 6 | 5BZO GSL |
| n115 | 684.1469 | 5.08 | 6'-sin-7MSOH GSL |
| n114 | 640.1211 | 7.61 | 6'-sin-5MTP GSL |
| n91 | 698.1627 | 5.6 | 6'-sin-8MSOO GSL form 1 |
| n167 | 654.1362 | 9.12 | 6'-sin-6MTH GSL |
| n185 | 668.152 | 11.63 | 6'-sin-7MTH GSL form 2 |
| n106 | 668.1522 | 10.71 | 6'-sin-7MTH GSL form 1 |
| n162 | 698.1628 | 6.12 | 6'-sin-8MSOO GSL form 2 |
| n139 | 642.1004 | 4.42 | 6'-sin-4MSOB GSL |
| n153 | 788.1582 | 6.29 | 6'-sin-3SIN GSL |
| n123 | 540.0682 | 5.21 | 6'-bnz-4-MSOB GSL |
| n17 | 448.0779 | 4.37 | 6-MTH GSL |
| n95 | 682.1677 | 11.96 | 6'-sin-8MTO GSL form 1 |
| n74 | 422.059 | 3.7 | Phenethyl GSL |
| n214 | 420.0468 | 2.47 | 4MTB GSL |
| n170 | 682.168 | 12.76 | 6'-sin-8MTO GSL form 2 |
| n14 | 422.026 | 0.68 | 3MSOP GSL |
| n158 | 524.0729 | 8.93 | 6'-sin-4MTB GSL |
| n111 | 540.0681 | 4.36 | 6'-sin-4MSOB GSL |
| n93 | 580.1357 | 13.92 | 6'-bnz-8MTO GSL |
| n103 | 566.1203 | 12.7 | 6'-bnz-7MTH GSL |
| n119 | 538.0889 | 9.88 | 6'-bnz-5MTP GSL |
| n96 | 551.0808 | 8.6 | 6'-bnz-indol-3-ylmethyl GSL |
| n156 | 552.1046 | 11.38 | 6'-bnz-6MTH GSL |
| n107 | 596.1309 | 6.73 | 6'-bnz-8MSOO GSL |

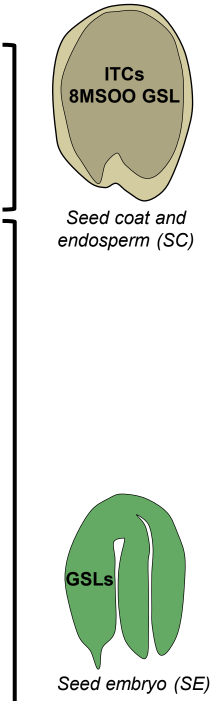


**Supplementary figure 7: Distribution of glucosinolate and related degradation products between seed coat & endosperm (SC) and seed embryo (SE) in *aop3-1* mutant and wild-type dry seeds developed under control and elevated temperature.** Metabolite identity (ID), mass/charge ratio (*m/z*), retention time (RT), putative annotation are listed for each glucosinolate and degradation product.

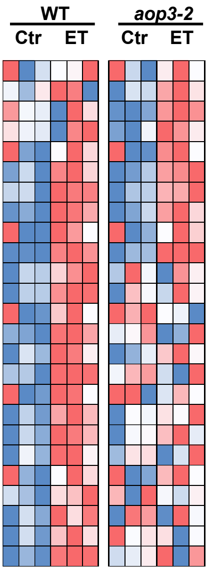


| Log$_2$(ET/Ctr) WT | Log$_2$(ET/Ctr) *aop3-2* | Average signal accumulation WT | Average signal accumulation *aop3-2* | ID | *m/z* | RT | Putative annotation | Spectral information |
|---|---|---|---|---|---|---|---|---|
| | 2.21 | *0.521* | *0.774* | n204 | 656.1174 | 4.59 | 6'-sin-5MSOP GSL | m/z 265; 447; 465 - losses 15,02; 63,99 |
| | 1.51 | *0.711* | *0.905* | n300 | 653.1127 | 7.40 | GSL_n300 | |
| | 2.23 | *5.553* | *15.757* | n45 | 450.0571 | 0.87 | 5MSOP GSL | m/z 259; 274 - losses 15,02; 63,99 |
| | 2.42 | *0.836* | *0.907* | p769 | 449.0682 | 3.47 | 3-Indolylmethyl GSL | |
| 2.18 | 1.60 | *170.200* | *232.865* | n152 | 390.0536 | 0.71 | Hydroxybutyl GSL isomer | |
| 2.06 | 1.16 | *1.186* | *1.681* | n182 | 540.0681 | 5.22 | 6'-bnz-4MSOB GSL | m/z 379; 395 |
| 3.05 | 1.41 | *0.453* | *0.334* | n390 | 552.0771 | 4.15 | GSL_n390 | |
| 2.36 | 1.18 | *11.572* | *18.591* | n46 | 406.0308 | 1.66 | 3MTP GSL | m/z 259; 274 |
| 2.38 | 1.64 | *1.068* | *0.699* | p528 | 174.0948 | 1.46 | GSL_p528 | |
| 6.07 | 7.40 | *0.092* | *0.051* | p749 | 480.1029 | 2.47 | 7MSOH | |
| 1.77 | | *1.276* | *1.738* | n147 | 524.0731 | 9.24 | 6'-bnz-4MTB GSL | m/z 363; 379; 395 |
| 2.05 | | *0.983* | *1.938* | n156 | 527.0732 | 8.47 | GSL_n156 | |
| 1.26 | | *25.224* | *33.075* | n19 | 422.0587 | 3.71 | 2-phenylethyl GSL | |
| 3.83 | | *0.489* | *1.272* | n214 | 422.0262 | 0.69 | 3MSOP GSL | m/z 259; 274 - losses 15,02; 63,99 |
| 1.53 | | *2.162* | *2.863* | n231 | 670.1314 | 4.75 | 6'-sin-6MSOH GSL | m/z 265; 447; 465 - losses 15,02; 63,99 |
| 5.87 | | *0.190* | *0.532* | n240 | 554.0836 | 4.62 | 6'-bnz-5MSOP GSL | m/z 363; 379; 395 - losses 15,02; 63,99 |
| 2.12 | | *1.609* | *2.696* | n245 | 582.1144 | 5.78 | 6'-bnz-7MSOH GSL | m/z 363; 379; 395 - losses 15,02; 63,99 |
| 1.71 | | *14.623* | *17.488* | n28 | 525.0682 | 4.29 | GSL_n28 | |
| 1.44 | | *1.363* | *1.621* | n281 | 653.1127 | 8.51 | GSL_n281 | |
| 1.31 | | *179.792* | *213.842* | n29 | 700.1383 | 10.07 | 6'-sin-4BZO GSL | m/z 465 |
| 1.76 | | *0.573* | *0.914* | n426 | 431.1927 | 3.97 | GSL_n426 | |
| 6.45 | | *0.022* | *0.024* | n454 | 508.0464 | 5.20 | GSL_n454 | |
| 1.81 | | *0.527* | *0.987* | n458 | 596.1304 | 7.05 | 6'-bnz-8MSOO GSL | m/z 363; 395 - losses 15,02; 63,99 |
| 1.14 | | *2.740* | *4.139* | n587 | 496.1006 | 3.52 | GSL_n587 | |
| 2.48 | | *0.755* | *0.889* | n98 | 460.0958 | 4.26 | GSL_n98 | |

*Minimal log$_2$(FC)* — *Maximal log$_2$(FC)*    *Minimal accumulation* — *Maximal accumulation*

**Supplementary figure 8: Glucosinolates affected by elevated temperature in *aop3-2* mutant and wild type at dry seed developmental stage.** Relative accumulation heatmaps, log$_2$(elevated temperature/control average total accumulation) (log$_2$[ET/Ctr]), control accumulation averages, metabolite identity (ID), mass/charge ratio (*m/z*), retention time (RT), putative annotation and spectral annotation are listed for each glucosinolate.

### Volcano plot  dry seed WT

144 induced SMs and 133 repressed SMs

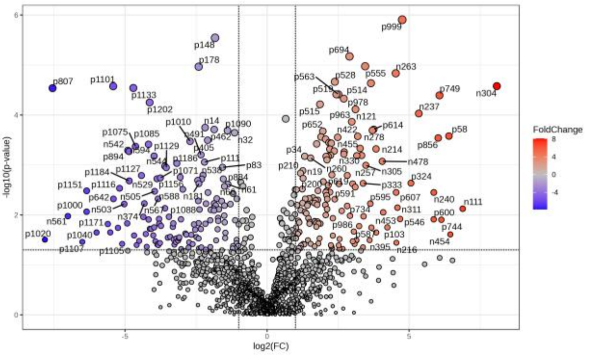


| | WT | |
|---|---|---|
| **Metabolic category** | **Induced** | **Repressed** |
| Flavonoids | 3 | 11 |
| Cinnamic acid and derivatives | 9 | 10 |
| Lipids and FA | 0 | 10 |
| Glucosinolates | 20 | 5 |
| Carbohydrates | 2 | 2 |
| Amino acids and derivatives | 2 | 1 |
| Choline derivatives | 1 | 1 |
| Glucosinolate degradation products | 1 | 1 |
| Benzaldehyde | 0 | 1 |
| Alkaloids | 0 | 1 |
| Indole derivative | 2 | 0 |
| Nucleosides and derivatives | 1 | 0 |
| Quinones | 1 | 0 |

### Volcano plot  dry seed *aop3-2*

78 induced SMs and  171 repressed SMs

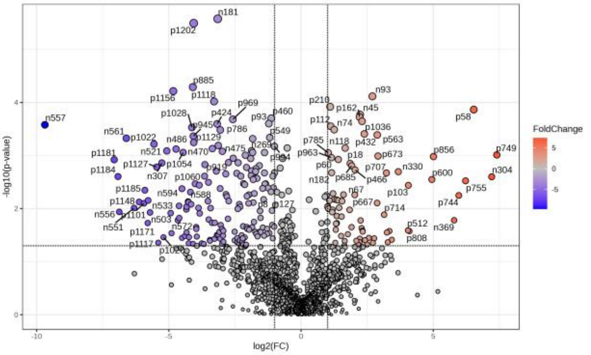


| | *aop3-2* | |
|---|---|---|
| **Metabolic category** | **Induced** | **Repressed** |
| Cinnamic acid and derivatives | 16 | 13 |
| Lipids and FA | 0 | 10 |
| Flavonoids | 1 | 10 |
| Glucosinolates | 9 | 6 |
| Benzaldehyde | 1 | 2 |
| Carbohydrates | 2 | 2 |
| Alkaloids | 0 | 1 |
| Benzoic acids | 0 | 1 |
| Carboxylic acids | 0 | 1 |
| Lignans | 0 | 1 |
| Terpenoids | 0 | 1 |
| Choline derivatives | 1 | 1 |
| Glucosinolate degradation products | 1 | 1 |
| Amino acids and derivatives | 5 | 1 |

**Supplementary figure 9: Volcano plot analyses performed to identify the specialized metabolites affected by elevated temperature in dry seeds of both *aop3-2* mutant and wild-type genotypes developed under control or elevated temperature.** Specialized metabolites with $\log_2$ [ET/Ctr] ≤ -1 and $\log_2$ [ET/Ctr] ≥ 1 were considered as repressed and induced by elevated temperature respectively. The number of induced and repressed metabolites belonging to known metabolic categories are given for each genotype and seed developmental stage.

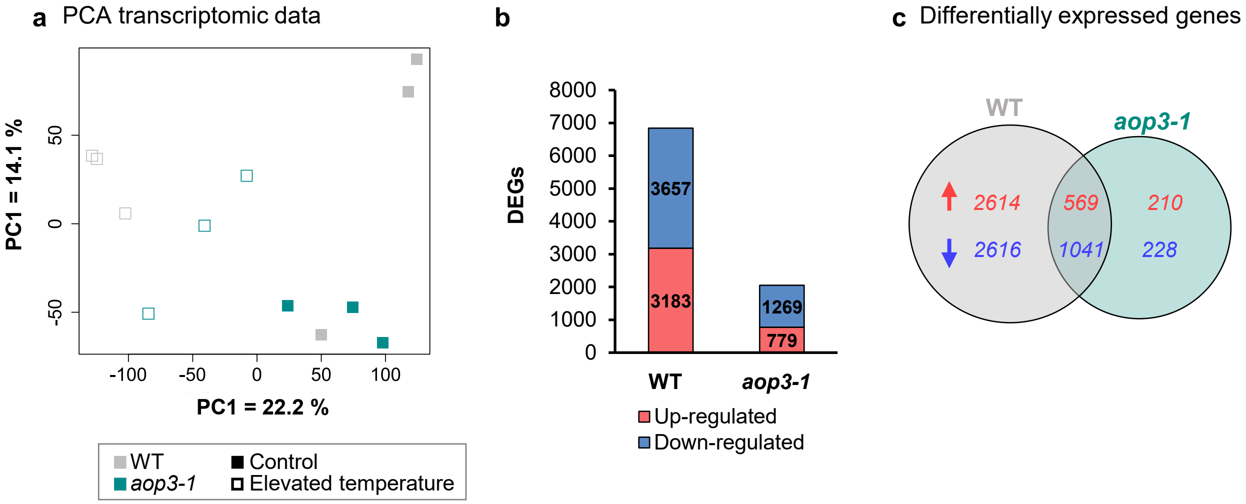


**Supplementary figure 10: Statistical analyses performed on transcriptomic data obtained from mature-green seed developmental stage of wild-type and *aop3-1* mutant genotypes developed under control or elevated temperature.** a) Principal component analysis. b) Genes differentially expressed (up or down-regulated) upon elevated temperature condition. c) Venn diagram presenting the genotype-specific and common up-regulated and down-regulated genes.

**a**

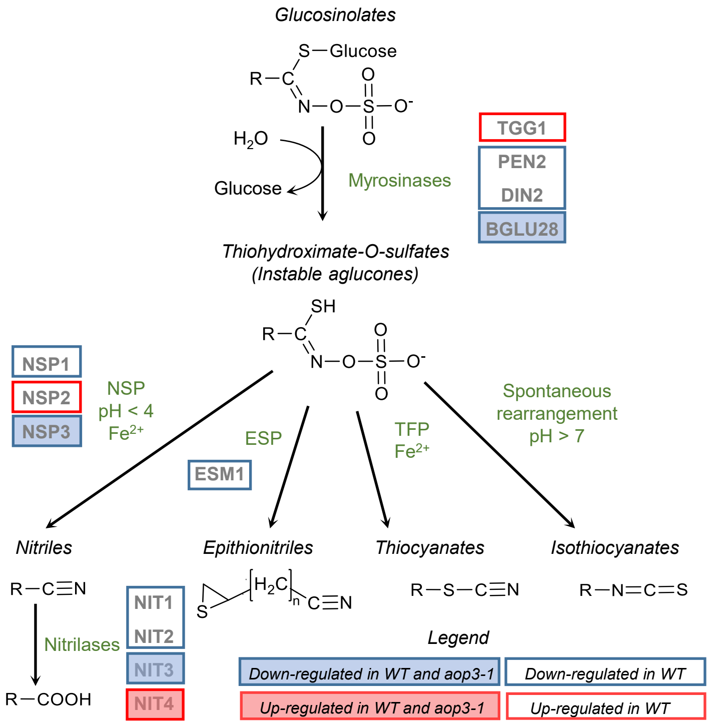


**DEGs involved in glucosinolate degradation**

| Gene name | AGI code | Acronym | WT | aop3-1 |
|---|---|---|---|---|
| epithiospecifier modifier 1 | AT3G14210 | ESM1 | x | |
| thigoglucosehydrolase 1 | AT5G26000 | TGG1 | x | |
| penetration 2 | AT2G44490 | PEN2 | x | |
| β-glucosidase 28 | AT2G44460 | BGLU28 | x | x |
| β-glucosidase 30 | AT3G60140 | DIN2 | x | |
| nitrilase 1 | AT3G44310 | NIT1 | x | |
| nitrilase 2 | AT3G44300 | NIT2 | x | |
| nitrilase 3 | AT3G44320 | NIT3 | x | x |
| nitrilase 4 | AT5G22300 | NIT4 | x | x |
| nitrile specifier protein 1 | AT3G16400 | NSP1 | x | |
| nitrile specifier protein 2 | AT2G33070 | NSP2 | x | |
| nitrile specifier protein 3 | AT3G16390 | NSP3 | x | x |

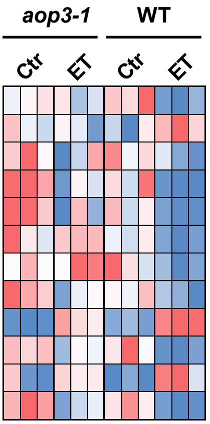


*Legend*
*x = gene differentially expressed*

**b**

**DEGs involved in glucosinolate transport**

| Gene name | AGI code | Acronym | WT | aop3-1 |
|---|---|---|---|---|
| glucosinolate transporter 2 | AT5G62680 | GTR2 | x | x |
| glucosinolate transporter 3 | AT1G18880 | GTR3 | x | |
| usually multiple acids move in and out transporters 29 | AT4G01430 | UMAMIT29 | x | x |
| usually multiple acids move in and out transporters 30 | AT4G01450 | UMAMIT30 | x | |

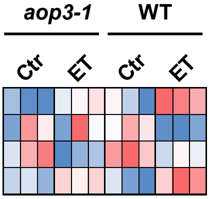


*Legend*
*x = gene differentially expressed*

**Supplementary figure 11: Effect of elevated temperature on genes coding for proteins involved in glucosinolate degradation and transport.** Heatmaps displaying the expression of differentially expressed genes coding for proteins involved in a) glucosinolate degradation and b) transport

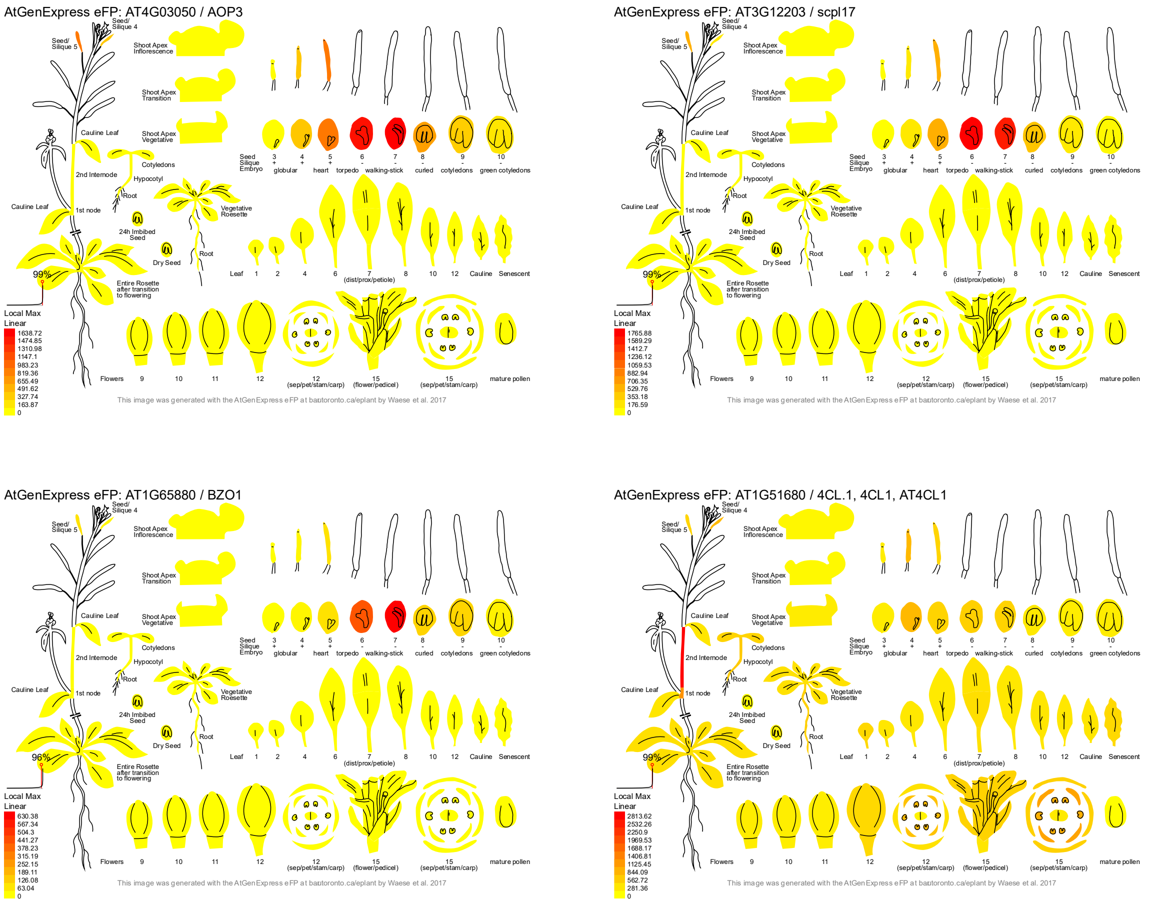


**Supplementary figure 12: *AOP3*, *SCPL17*, *BZO1* and *4CL* gene expression atlas**. Data from https://bar.utoronto.ca/eplant (Klepikova et al., 2016).

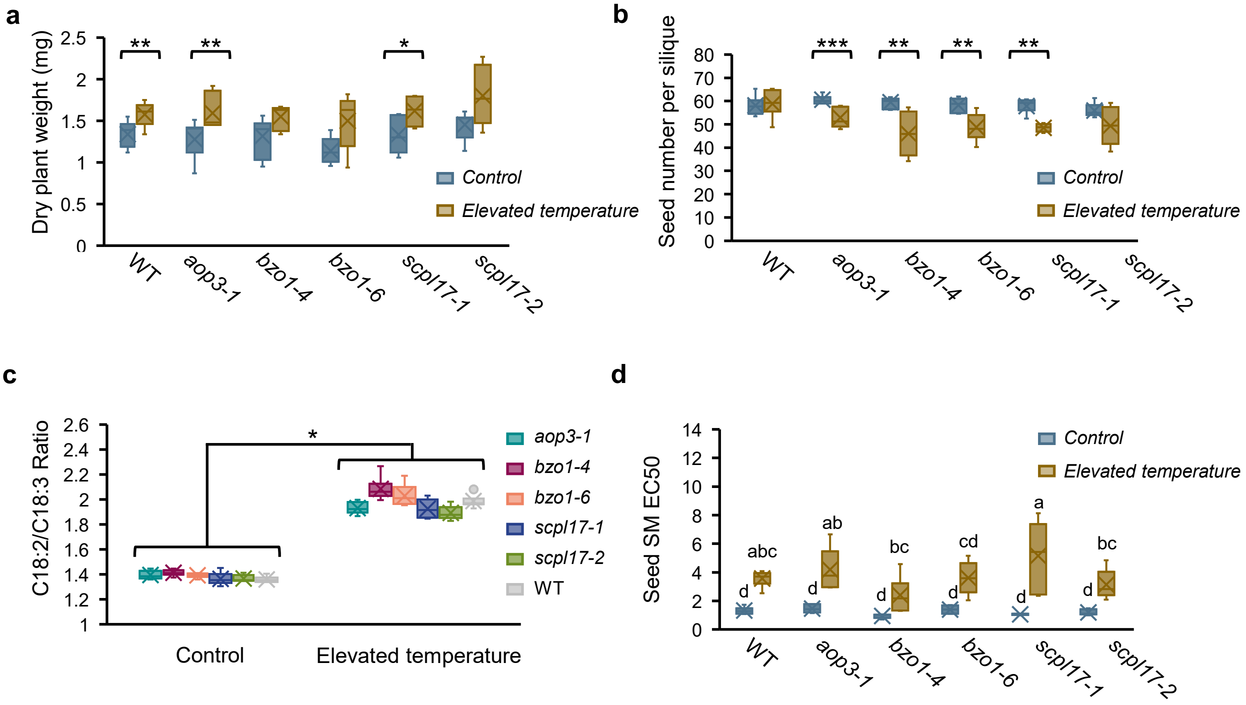


**Supplementary figure 13: Phenotypic analysis of *aop3*, *bzo1* and *scpl17* and wild-type plants and seeds developed under control and elevated temperature conditions.** a) Dry plant weight of *aop3*, *bzo1* and *scpl17* mutants and wild-type plants developed under control and elevated temperature conditions. Differences among conditions are indicated (Mann-Whitney tests - p-value ≤ 0.05 [*], p-value ≤ 0.01[**]) b) Number of seed per silique for *aop3*, *bzo1*, *scpl17* and wild-type genotypes grown under control and elevated temperature conditions. Differences among conditions are indicated (Mann-Whitney tests - p-value ≤ 0.01 [**], p-value ≤ 0.001[***]) c) C18:2/C18:3 ratio in dry seeds of *aop3*, *bzo1*, *scpl17* and wild-type lines grown under control and elevated temperature conditions. Differences among conditions are indicated (Kruskal-Wallis test - p-value ≤ 0.05 [*]. d) Antioxidant capacities of polar and semi-polar metabolites extracted from dry seeds of *aop3*, *bzo1*, *scpl17* and wild-type genotypes developed under control and elevated temperature conditions. The smallest de EC50 value (effective concentration which scavenges 50% of DPPH [2,2-diphenyl-1-picrylhydrazyl] radical) is, the strongest is the antioxidant capacity. Differences among conditions are indicated (ANOVA test, Newman-Keuls post-hoc test).

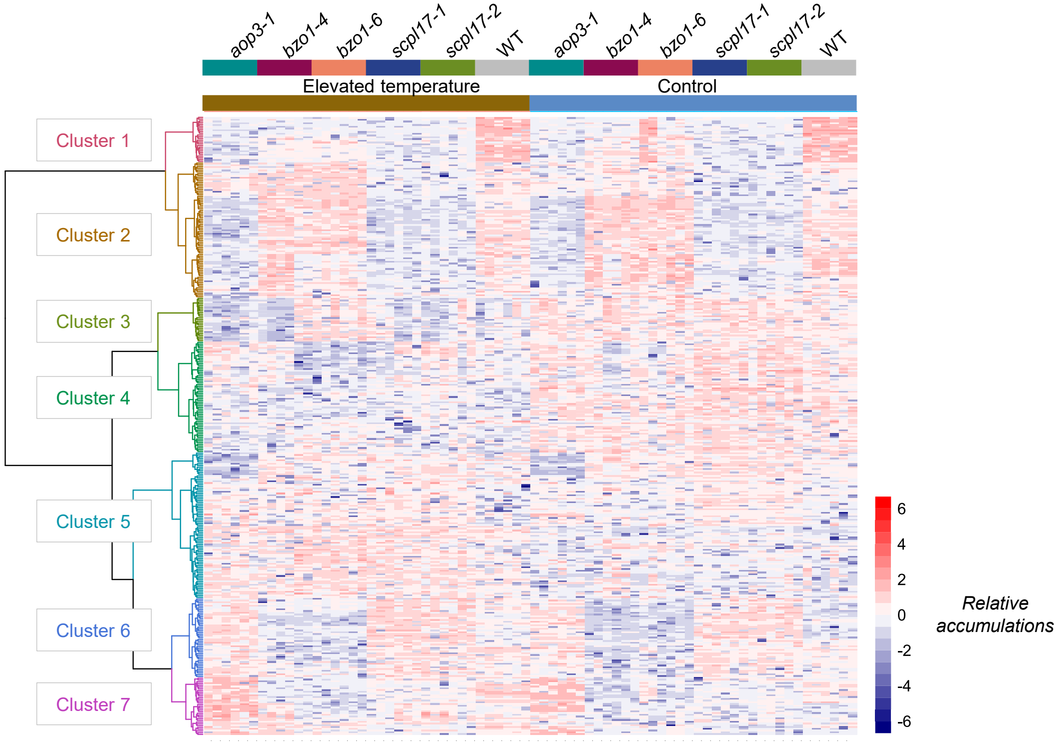


| Cluster 1 | |
|---|---|
| Cinnamic acid and derivatives | 4 |
| Glucosinolates | 3 |

| Cluster 2 | |
|---|---|
| Benzaldehyde | 1 |
| Carboxylic acid | 1 |
| Cinnamic acid and derivatives | 9 |
| Flavonoids | 1 |
| Glucosinolates | 11 |
| Lignans | 1 |
| Lipids and FA | 1 |

| Cluster 3 | |
|---|---|
| Alkaloids | 2 |
| Flavonoids | 2 |
| Lipids and FA | 12 |

| Cluster 4 | |
|---|---|
| Cinnamic acid and derivatives | 10 |
| Flavonoids | 4 |
| Glucosinolates | 2 |
| Glucosinolate degradation products | 3 |
| Lipids and FA | 5 |
| Nucleosides | 1 |

| Cluster 5 | |
|---|---|
| Benzaldehyde | 1 |
| Benzoic acid | 1 |
| Carbohydrates | 2 |
| Cinnamic acid and derivatives | 7 |
| Flavonoids | 9 |
| Glucosinolates | 9 |
| Lipids and FA | 1 |

| Cluster 6 | |
|---|---|
| Carbohydrates | 2 |
| Cinnamic acid and derivatives | 8 |
| Flavonoids | 2 |
| Glucosinolates | 3 |

| Cluster 7 | |
|---|---|
| Amino acids and derivatives | 2 |
| Cinnamic acid and derivatives | 1 |
| Flavonoids | 1 |
| Glucosinolates | 6 |

**Supplementary figure 14: Heatmap displaying the relative accumulation of the specialized metabolites that were determined to be affected by elevated temperature and/or genotype.** The number of metabolites belonging to known metabolic categories are given for each heatmap cluster (defined according to specialized metabolite accumulation profiles).

**Log$_2$(Mutant/WT control accumulations)**

| *aop3-1* | *bzo1-4* | *bzo1-6* | *scpl17-1* | *scpl17-2* | | *m/z* | RT | Putative annotation | Cluster heatmap |
|---|---|---|---|---|---|---|---|---|---|
| -3.43 | -3.46 | -2.16 | -2.83 | -3.23 | n128 | 584.0919 | 10.81 | 6'-bnz-3BZO GSL | 1 |
| -2.89 | -1.36 | -1.20 | -2.92 | -2.49 | n26 | 549.0860 | 3.87 | GSL_n26 | 1 |
| -2.95 | -0.77 | -0.07 | -3.73 | -3.61 | n35 | 700.1384 | 9.90 | 6'-sin-4BZO GSL form 1 | 1 |
| -2.22 | 1.65 | 1.16 | -1.52 | -0.93 | n146 | 802.1700 | 8.23 | 6'-sin-4SIN GSL | 2 |
| -0.91 | -0.37 | -0.52 | -0.91 | 0.23 | n15 | 390.0537 | 0.74 | Hydroxybutyl GSL isomer | 2 |
| -1.67 | 2.50 | 2.33 | -1.36 | -1.26 | n203 | 788.1543 | 7.22 | 6'-sin-3SIN GSL | 2 |
| -0.03 | -0.22 | 0.55 | -1.18 | -1.00 | n213 | 654.1364 | 9.20 | 6'-sin-6MTH GSL | 2 |
| -2.42 | -0.47 | -0.58 | -2.16 | -1.58 | n24 | 563.1017 | 4.14 | GSL_n24 | 2 |
| -1.48 | 1.22 | 1.73 | -1.44 | -1.81 | n290 | 610.1281 | 5.64 | GSL_n290 | 2 |
| -1.90 | 0.02 | 0.43 | -2.70 | -2.34 | n46 | 686.1228 | 8.66 | 6'-sin-3BZO GSL | 2 |
| -3.31 | 0.32 | 0.05 | -3.69 | -3.38 | n54 | 510.0752 | 4.02 | GSL_n54 | 2 |
| 0.26 | 0.28 | 0.36 | -1.42 | -1.43 | n71 | 640.1206 | 7.67 | 6'-sin-5MTP GSL | 2 |
| -2.88 | -1.79 | -0.73 | -3.02 | -2.86 | n84 | 700.1385 | 10.81 | 6'-sin-4BZO GSL form 2 | 2 |
| -2.31 | -0.56 | -0.01 | -1.09 | -1.35 | n94 | 582.0965 | 4.45 | 3SIN GSL | 2 |
| -1.63 | 0.21 | 0.07 | 0.77 | 0.34 | n201 | 418.0313 | 2.97 | GSL_n201 | 4 |
| -1.49 | -1.86 | -1.74 | -1.79 | -1.73 | n62 | 508.0962 | 6.05 | 5BZO GSL | 4 |
| -0.61 | -0.68 | -0.04 | 0.54 | 0.40 | p483 | 220.0823 | 21.47 | 7MSOH ITC | 4 |
| -0.66 | -1.50 | -0.16 | 0.28 | 0.40 | p509 | 218.1036 | 22.54 | ITC_p509 | 4 |
| -0.03 | -1.20 | 0.37 | 1.07 | 0.34 | p524 | 170.0994 | 22.54 | ITC_p524 | 4 |
| 2.23 | -0.29 | -0.48 | 0.26 | 0.14 | n131 | 510.0570 | 6.78 | GSL_n131 | 5 |
| -2.64 | -0.07 | 0.34 | 0.62 | 0.62 | n205 | 372.0434 | 1.24 | 3BUT GSL | 5 |
| -2.16 | 0.37 | -0.23 | 0.62 | -0.24 | n227 | 596.1118 | 4.48 | GSL_227 | 5 |
| -0.84 | 1.03 | 1.23 | 0.98 | 0.69 | n25 | 596.1119 | 4.92 | GSL_n25 | 5 |
| -3.25 | 0.34 | 0.79 | 1.32 | 1.20 | n286 | 404.0696 | 0.86 | GSL_n286 | 5 |
| -1.68 | -0.90 | 0.57 | 1.04 | 0.35 | n29 | 495.0756 | 2.59 | GSL_n29 | 5 |
| 0.44 | 1.16 | 0.10 | 1.52 | 1.81 | n63 | 682.1677 | 12.02 | 6'-sin-8MTO GSL | 5 |
| -1.31 | 0.81 | 1.02 | 1.04 | 0.98 | n76 | 376.0380 | 0.67 | GSL_n76 | 5 |
| 0.79 | 1.42 | 1.19 | -0.06 | -0.26 | n82 | 480.1054 | 7.13 | GSL_n82 | 5 |
| 1.41 | -2.32 | -0.51 | 0.57 | 0.24 | n137 | 580.1359 | 13.97 | 6'-bnz-8MTO GSL | 6 |
| 1.71 | -2.15 | -0.68 | 0.01 | -0.44 | n157 | 596.1307 | 6.80 | 6'-bnz-8MSOO GSL | 6 |
| -0.11 | -1.24 | -0.45 | 0.82 | 1.05 | n19 | 524.0731 | 8.19 | 6'-bnz-4MTB GSL form 1 | 6 |
| 1.48 | -2.55 | -1.22 | -2.12 | -1.66 | n115 | 551.0806 | 8.66 | 6'-bnz-indol-3-ymethyl GSL | 7 |
| 1.11 | -0.99 | -0.16 | -1.82 | -2.36 | n123 | 540.0682 | 5.23 | 6'-bnz-4MSOB GSL form 2 | 7 |
| 3.10 | 0.79 | 1.36 | 0.00 | 0.58 | n132 | 612.0892 | 5.89 | GSL_n132 | 7 |
| 1.94 | -1.82 | 0.03 | -0.09 | -0.48 | n142 | 540.0681 | 4.38 | 6'-bnz-4MSOB GSL form 1 | 7 |
| 2.19 | -1.89 | -0.34 | -0.25 | -0.54 | n165 | 538.0888 | 9.94 | 6'-bnz-5MTP GSL | 7 |
| 2.17 | -1.60 | -0.82 | -1.12 | -0.42 | n209 | 524.0729 | 9.00 | 6'-bnz-4MTB GSL form 2 | 7 |

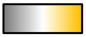


**Supplementary figure 15: Differences of glucosinolate accumulations in *aop3*, *scpl17* and *bzo1* mutant compared to wild type in dry seeds**. The log$_2$ (Mutant/WT control accumulations), metabolite identity (ID), mass/charge ratio (*m/z*), retention time (RT) and putative annotation are listed for each glucosinolate.
